# Reconstructing Actin Dynamics of the Leading Edge from Observational Data

**DOI:** 10.64898/2026.01.31.703047

**Authors:** Wenzheng Shi, Christopher E. Miles, Jungsik Noh, Gaudenz Danuser, Alex Mogilner

## Abstract

Mesenchymal cell migration starts with the protrusion of the cell’s leading edge, enabled by the branching growth of the actin network. Two crucial molecular players in protrusion are actin filaments and the Arp2/3 protein complex, the branching agent. Traditionally, protrusion models are intuited from several perturbative experiments. Recent multiplex microscopy imaging data promise to drastically change this paradigm by providing measurements of naturally fluctuating densities of key proteins at the leading edge of unperturbed cells. We report the first attempt to reconstruct the mechanistic protrusion model from such data. We analyze fluctuations of F-actin and Arp2/3 densities and cell edge velocity using phase space and regression analysis to reconstruct linearized stochastic actin-Arp2/3-velocity coupled dynamics. We then build a nonlinear partial differential equation model of these dynamics based on previous knowledge of this system and parsimony. The resulting model recovers the rates of reactions and mechanical and transport processes in the lamellipodium from a single non-perturbative experiment. The model posits that the only essential nonlinearities in the lamellipodial dynamics stem from the retrograde flow of F-actin. The model suggests that the protrusion is dominated by a damped oscillatory cycle resulting from a combination of positive feedback from Arp2/3 to protrusion velocity and negative feedback from F-actin to the velocity.

**Significance statement:** Traditionally, models of intracellular mechanochemistry are intuited from perturbative experiments. Multiplex microscopy imaging changes this by providing measurements of naturally fluctuating densities of key proteins in unperturbed cells. We use such data to reconstruct a mechanistic model of cell leading edge. We build equations for actin system and edge velocity using data, previous knowledge and parsimony. The resulting model recovers the rates of reactions and mechanical and transport processes at the leading edge. The model suggests that the protrusion is dominated by a damped oscillatory cycle resulting from a combination of positive feedback from Arp2/3 to protrusion velocity and negative feedback from F-actin to the velocity.

## INTRODUCTION

Deciphering the protruding leading edge machinery during mesenchymal cell migration has attracted much effort from cell biologists [1]. The protrusion is driven by a dynamic branched actin network of the motile appendage called lamellipodium [1–6]. The lamellipodial dynamics are captured by the dendritic nucleation model [2], according to which: 1) At the very edge, nucleation promoting factors, such as WAVE [7–10], activate Arp2/3 complex (Arp2/3 for brevity from here on). 2) Arp2/3 mediates branching of actin filaments (F-actin), growing plus ends of which push on the membrane, generating protrusion [6, 11]. 3) The filaments’ plus ends are capped [12], and then the filaments are disassembled [13], while the actin subunits and accessory molecules get recycled to the edge [14]. These dynamics are organized by dozens of actin-binding proteins with distinct structural and kinetic properties [1, 5, 15]: besides the branching agent Arp2/3 and capping protein, there are plus-end nucleators and elongators formin and VASP, disassembly governor cofilin and many more [12, 13, 15–20].

The flat (about 0.1 micron-thin [4]) lamellipodial actin network, adhering to the substrate, occupies the 1–2 microns wide zone at the leading edge [9, 21], bordered at the rear by a thicker, more bundled and contractile, lamellar actomyosin network [21, 22]. Lamellipodial mechanics stem from the balance of several principal forces: Actin filaments, growing with rates on the order of a micron per minute [21], push on the edge, generating membrane tension [23]. This tension both slows down the filament growth [11, 23, 24] and pushes the whole actin network back [25]. This push, assisted by a myosin-powered pull from the lamellipodium-lamellum boundary, causes the network to flow retrogradely (with rates on the order of a micron per minute [21]). The retrograde flow rate is determined by the balance between the membrane pushing plus myosin pulling force and the adhesion-based resistance to the flow [26–31].

Much of our knowledge about the cell edge came from the reductionist agenda approach [4] based on breaking complex dynamics into mechanochemical modules and studying them in isolation by perturbation experiments. Modeling was an integral part of this agenda [4, 32]. Both continuous deterministic differential-equation models [33–39] and discrete stochastic agent based models [11, 40–45], and hybrid continuous-discrete models [46] were used to reproduce observed movements and protein distributions at the protruding edge and to decipher mechanochemical feedbacks as well as related adhesion and myosin dynamics [31, 47, 48].

Lessons from perturbation experiments and accompanying traditional modeling are limited by complex and redundant connections between mechanochemical pathways; inferring feedbacks between these pathways from perturbed lamellipodial states is often impossible [49]. One of the ways to approach this conundrum is to observe natural molecular fluctuations in the unperturbed system and to deduce molecular feedbacks from the measured fluctuations [49, 50]. However, statistical cross-correlations between fluctuating protein densities offer very limited insight [49, 50]. Analysis of causality in time-resolved fluctuation data — test of whether past fluctuations of one protein signal contribute statistically significantly to prediction of a current concentration of another protein — is a more powerful approach. Recent application of Granger Causality to multiplex data from the lamellipodial edge [51] suggested that there are feedbacks between Arp2/3 and F-actin, and between F-actin and protrusion velocity, but no direct feedbacks between Arp2/3 concentration and velocity. Causality approaches do not provide explicit mechanistic description of the system. In comparison, the dynamical systems approach offers an intuitive understanding by describing the system in terms of differential equations, in addition to estimates of the relative strengths of its feedbacks [50, 52].

There has been a recent surge in combining applied math and data science to develop frameworks for reconstructing models from data [52–55]. A tool that leverages spectral basis representations and sparse regression algorithms to discover partial differential equations (PDE) models from microscopic simulation and experimental data was described in [56]. The Sparse Identification of Nonlinear Dynamics (SINDy) approach combines sparsity-promoting techniques and machine learning with nonlinear dynamical systems and allows discovering equations from noisy data [57]. Replacing derivative approximations with linear transformations and variance reduction techniques allows one to tackle systems with large noise [58]. Physics-informed neural networks with sparse regression were applied to discover PDEs from scarce and noisy data [59]. Identifying stochastic differential equations (SDEs) using sparse Bayesian learning was introduced in [60]. Experimental noise poses a challenge for these approaches that more often than not were used to analyze synthetic rather than experimental data [52].

In this study, we reconstruct a predictive mechanistic model of interactions between Arp2/3, F-actin, and cell edge movements from high-resolution observational data reported in [51]. We begin by using time series for the two protein concentrations and edge velocity to build a multidimensional phase portrait of the system. This results in a linear dynamical system whose coefficients are learned from the data. By applying sparsity and parsimony arguments, together with prior biophysical knowledge, we construct a nonlinear system of stochastic partial differential equations capturing spatial-temporal feedbacks that govern edge dynamics.

The model suggests that the protrusion velocity is determined by the ratio of the resisting force, proportional to the global F-actin length, to the density of edge-pushing actin barbed ends, proportional to the Arp2/3 density at the front. The model posits that the lamellipodial dynamics are dominated by the following cycle: Arp2/3 is integrated into the actin network mostly at the leading edge; filaments branched from these Arp2/3 grow and push the edge forward. Higher protrusion velocity makes these filaments treadmill away and spread over the lamellipodial width faster. Filaments’ attachments to adhesion complexes and membrane generate mechanical resistance, slowing the protrusion down. The model provides estimates of Arp2/3-actin-velocity dynamic parameters simply from measuring mechanochemical fluctuations in an unperturbed lamellipodium.

## RESULTS

### Phase-space data analysis reconstructs linearized lamellipodial dynamics

Tracking dynamic leading edge [61] of the narrow lamellipodium (∼ 1.5 *µ*m wide) provides the data for our modeling effort (Fig. 1A, SI Appendix). The lamellipodium is discretized into the approximately rectangular grid (Fig. 1B) with two rows of ≈ 0.7 *µ*m × 0.7 *µ*m square compartments, the size of which is limited by the microscopy resolution [51]. The leading compartments border the lamellipodial edge. Local edge protrusions are tracked every 3 sec for 15 min, and locally normal velocities of the leading compartment edges are measured. The grid is shifting and deforming in the lab frame together with the fluctuating leading edge, and so in the moving frame of the cell edge, the compartments preserve their local positions relative to the edge. The sampling of time-lapse images of U2OS cells co-expressing fluorescent Arp2/3 and actin in these deformable grids provide us with time series for fluctuating Arp2/3 and F-actin densities in each of the lamellipodial compartments [51]. We normalize (cell by cell) the time series for the velocity and Arp2/3 and F-actin densities by scaling the standard deviations of each of these three signals to unity and shifting respective means to zero (see Methods and SI Appendix; below we compare results from individual cells to those obtained from pooling all cells).

**FIG. 1.**
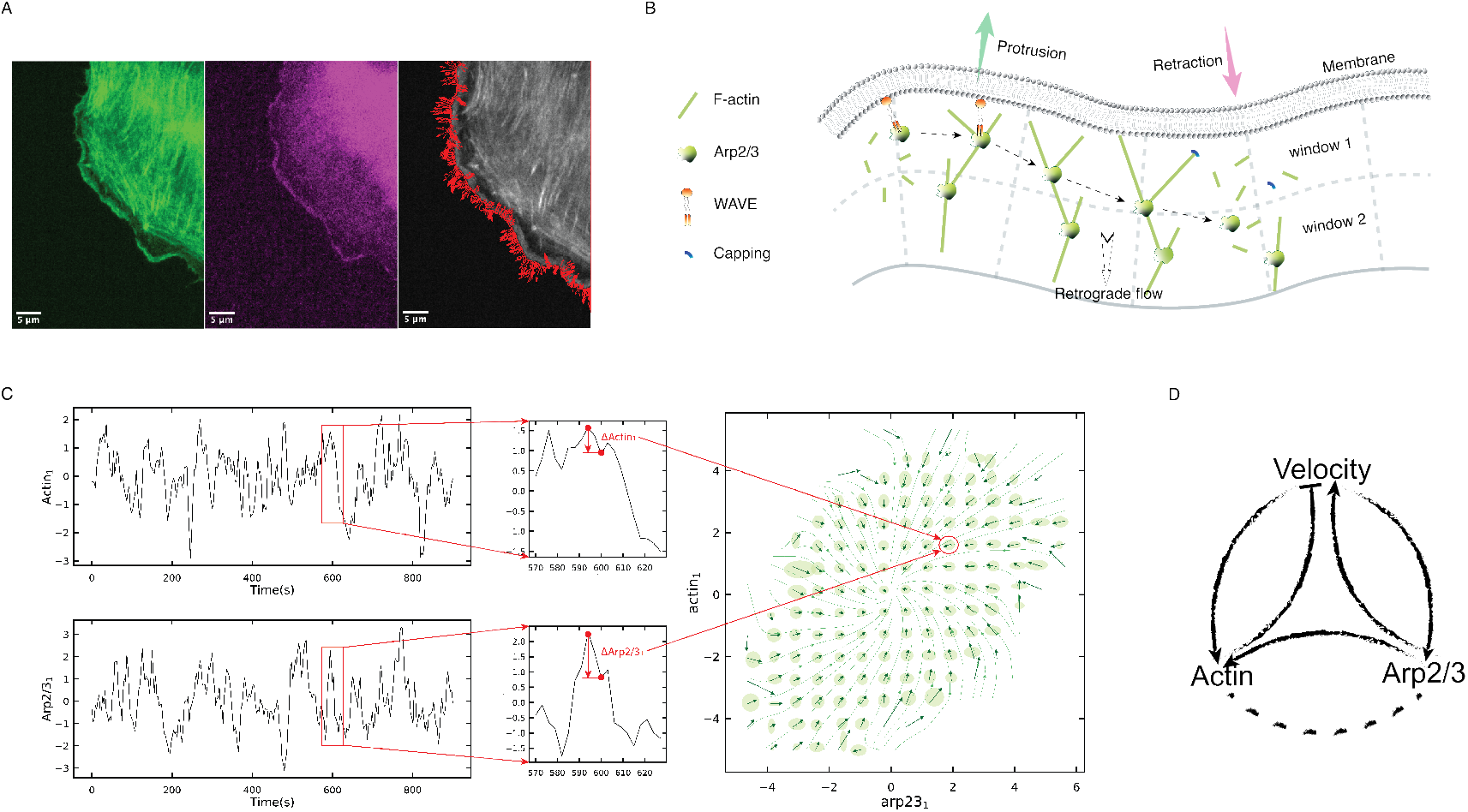
Phase-space analysis of velocity=actin-Arp2/3 dynamics. (A) Sample images on which the analysis is based are from live U2OS cells expressing fluorescent markers for Actin and Arp2/3. Left: F-actin, center: Arp2/3. Lamellipodia are seen as bright rims at the cell’s leading edge. Right: edge velocity (red arrows) is measured using a window-tracking algorithm. Further details are in Methods and in [51]. Scale bars: 5*µ*m. (B) Schematic of the branched actin network at the cell edge. Principal proteins, Arp2/3, WAVE, and capping protein, together with actin filaments, key movements, and the grid of leading (window 1) and trailing (window 2) compartments are shown. (C) Dynamic coupling between F-actin and Arp2/3 revealed through phase-space analysis. Left: time series of F-actin (top) and Arp2/3 (bottom) fluorescent signals from a representative probing window. Middle: zoomed-in region highlighting signal increments over a small time interval. Right: ensemble-averaged direction field in the phase space of Arp2*/*3_1_ (x-axis) and F − actin_1_ (y-axis). Arrows indicate the average direction of change in each bin, and green ellipses denote the uncertainty (standard deviation) in each direction. The data is from 12 cells, with total 972 windows. In each window, 15 min long time series with a time step d*t* = 3*s* is analyzed. (D) The feedback diagram (based on Eq. (1)) illustrates a preliminary Arp2/3-actin-velocity feedback network in a leading window. Arrows/bars indicate ‘excitation/inhibition’, respectively. The dashed line indicates the tentative character of the feedback from actin to Arp2/3.

We used the phase portrait analysis as a starting point for lamellipodial dynamics reconstruction. As an example, let us consider the problem of reconstructing just actin-Arp2/3 dynamics in one of the lamellipodial compartments, without analyzing the velocity effects. Let *A*(*t*) and *R*(*t*) be the time series for the F-actin and Arp2/3 data, respectively. At each time step, we assign a vector [(*A*(*t* + *dt*) − *A*(*t*)), (*R*(*t* + *dt*) − *R*(*t*))] to a point with coordinates [*A*(*t*), *R*(*t*)] on the phase plane, where *dt* = 3 sec is the time increment (Fig. 1C). Such vector shows the evolution of the actin-Arp2/3 system over a short time interval. Repeating this procedure for each time point results in the phase portrait (Fig. 1C). Of course, due to stochastic effects, several different vectors correspond to some closely located points on the *A* − *R* plane, so we discretize the phase plane with a grid, and for each grid point find the mean of all vectors closest to this point as well as the vectorial noise (Fig. 1C, Methods and SI Appendix).

This results in the phase portrait revealing that the system is characterized by a stable spiral (Fig. 1C, Fig. S1), so the actin-Arp2/3 densities are returning to their average values after being perturbed by the noise, the nature of which we discuss below. This phase portrait corresponds to a dynamical system of general form: *dA/dt* = *f* (*A, R*), *dR/dt* = *g*(*A, R*). For a leading-edge compartment, we expect the boundary velocity to be an intimate part of the dynamics, and the phase portrait analysis of the three time series, for actin, Arp2/3, and velocity (*V* ), gives the 3D phase portrait (Fig. S2). We used linear regression to find the least square error approximations of the linear dynamical *v* − *a* − *r* system (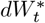 are the increments of the stochastic Wiener process; we use lower case *v* − *a* − *r* notations for the linear system and upper case *V* − *A* − *R* for the full nonlinear system, more below):

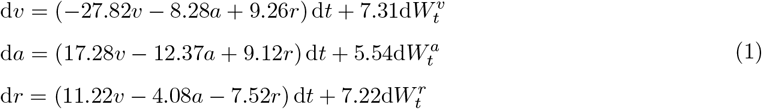

One of the mathematical terms in the right-hand-side of system Eq. (1) – (−4.08*a*) in the third equation – is relatively small, and one open question is whether there is enough confidence that this term is significant. We used sparsity arguments by employing an information criterion assigning a weight to the number of model parameters, in addition to the approximation error, and minimizing the combined score of the model. Specifically, we used LASSO regression (Fig. S3, SI Appendix) to screen the terms of system Eq. (1) and found that all terms are meaningful.

The reconstructed system Eq. (1) posits the following tentative dynamic links at the leading edge (Fig. 1D): 1) There are positive feedbacks from Arp2/3 to F-actin and velocity, which are easy to explain: the branching action of Arp2/3 generates new filaments near the leading edge, which push the membrane forward, accelerating the edge. 2) There is a negative feedback from F-actin to velocity and a tentative negative feedback from F-actin to Arp2/3 (corresponding to the smallest term in Eq. (1); this feedback is absent in the final model, below). Interpretation of these links is an open question. It is important to avoid the following potential confusion: actin filaments push the cell membrane at the leading edge forward, but the pushing force is proportional to the *number* (not to the total *length*) of these active leading filaments, and so to the number of Arp2/3 complexes at the front. On the other hand, the F-actin density feeding back negatively to the velocity is the *length* density of the filaments. 3) There are positive feedbacks from velocity to Arp2/3 and F-actin. Interpretation of these links is an open question. 4) The large negative diagonal terms (*dr/dt* ∼ −*r* etc.) can be tentatively interpreted as follows: for Arp2/3 and actin, they are due both to network disassembly and to transport away from the leading edge by the retrograde flow. For velocity, the term *dv/dt* ∼ −*v* can be interpreted as relaxation of the protrusion velocity to the level set by the Arp2/3 and F-actin densities.

Before we address these open questions, we have to investigate the interdependencies of the dynamic variables in the neighboring windows. First, the retrograde flow of the actin network, in the coordinate system moving with the cell edge, generates transport between the two rows of the lamellipodial compartments. Coupling the dynamics of velocity and of F-actin and Arp2/3 densities in the leading (*R*_1_ and *A*_1_) and trailing (*R*_2_ and *A*_2_) compartments gives us a 5-dimensional dynamical system of the form: *dV/dt* = *h*(*V, A*_1_, *R*_1_, *A*_2_, *R*_2_), *dA*_1,2_*/dt* = *f*_1,2_(*V, A*_1_, *R*_1_, *A*_2_, *R*_2_), *dR*_1,2_*/dt* = *g*_1,2_(*V, A*_1_, *R*_1_, *A*_2_, *R*_2_). Second, there can be physical transport between any lamellipodial compartment and neighboring compartments at the right and left. We found that respective terms are well approximated by effective diffusive transport between neighboring windows.

In this biologically inspired dynamics reconstruction, from the beginning, we omit terms for which we would not be able to find biophysical interpretation. Those include: 1) *r*_2_ term in the equation for *dv/dt*: it would be hard to explain how Arp2/3 at the lamellipodial rear affects the edge velocity. 2) *r*_2_ and *a*_2_ terms in the equations for *da*_1_*/dt* and *dr*_1_*/dt*: it would be hard to explain how Arp2/3 and F-actin at the lamellipodial rear affect Arp2/3-F-actin dynamics at the front, considering that characteristic filament size is smaller than the compartment size and that actin network flows from the front to the rear in the framework of the cell edge. We considered potential complex effects of limited conserved molecular resources in the lamellipodium shared between the network and cytoplasm and concluded that such effects would not explain the excluded terms without significantly worsening reconstruction of the whole system. 3) *r*_1_ term in the equation for *da*_2_*/dt* and *a*_1_ term in the equations for *dr*_2_*/dt*: it would be hard to explain such non-local influences.

Thus, we used the linear regression to find the least square error linear approximation of the spatial-temporal lamellipodial dynamics by assuming that these seven terms are absent. The resulting stochastic dynamical system, the full 5-variable system for the leading/trailing pair of windows (with spatial diffusion terms shown to just to illustrate lateral transport that will be properly addressed in the PDE model below), is shown in Fig. 2C. We found eigenvalues of the matrix of the corresponding deterministic linear system, all of which had negative real parts indicating stability of the steady state. Several eigenvalues had imaginary parts suggesting that the system is characterized by damped oscillations.

**FIG. 2.**
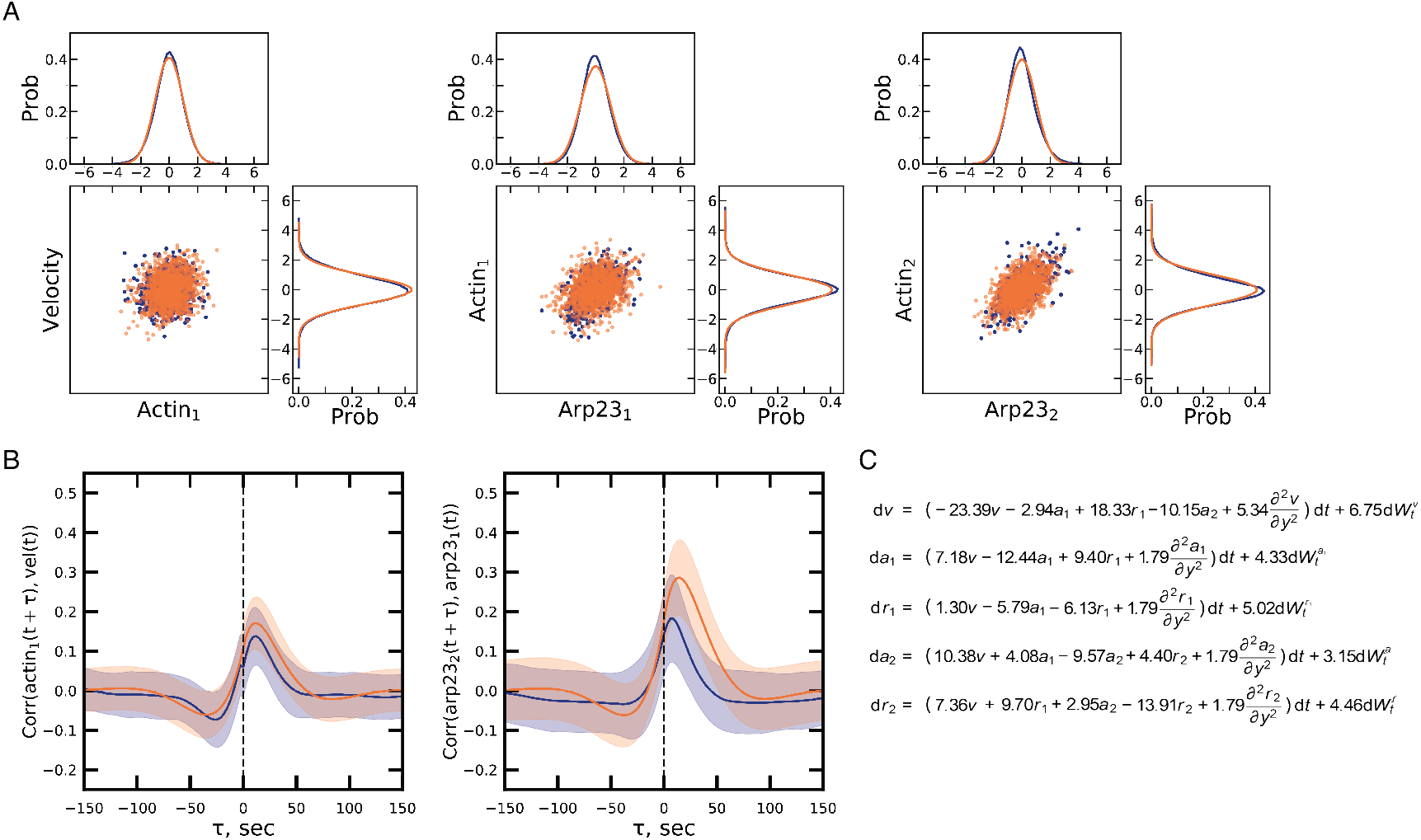
Statistics and correlations in lamellipodia are captured by the reconstructed linear model. (A) Comparison of experimental data (blue) and model simulations (orange) for the distributions of F-actin, Arp2/3, and velocity. Joint 2D scatter plots (bottom left of each panel) reveal elliptical distributions indicative of strong correlations between components, while marginal 1D distributions (top and right) confirm approximately Gaussian statistics for each variable, consistent with linear stochastic dynamics. (B) Ensemble-averaged cross-correlation functions between F-actin in leading compartments and velocity (left), and between Arp2/3 in leading and trailing compartments (right). Experimental curves are in blue, model results are in orange. The vertical dashed line denotes zero lag, *τ* = 0. Shaded bands mark *±* standard deviation across window ensembles. The data in (A) and (B) is from 12 cells, with total 972 windows. In each window, 15 min long time series with a time step d*t* = 3*s* is analyzed. (C) Final, reduced system of linear stochastic partial differential equations inferred from the data (see the text for explanation of the notations and terms).

#### Several notes on the dynamical system reconstruction: 1)

We used linear and non-linear regressions to find the least square error approximations of functions *h, f*_1,2_, *g*_1,2_ and found that non-linear terms are small and do not improve the model accuracy (SI Appendix). This indicates that though the system is inherently nonlinear [33, 34], the noise is not too great to perturb the dynamic equilibrium enough to reveal these nonlinearities. Only linearized dynamics close to the equilibrium can be deduced from the data. A similar conclusion was reached in [51].

2) When we compute the 2 × 2 matrix of the linear coefficients in the reconstructed *a* − *r* dynamical system for a single compartment and compare the matrix elements with the respective 2 × 2 sub-matrix of the 3 × 3 matrix of the reconstructed *v* − *a* − *r* system for the single compartment, the respective elements remain very close to each other. Similarly, 9 elements of the 3 × 3 matrix of the reconstructed *v* − *a* − *r* system for the single compartment are close to respective coefficients of the *v* − *a*_1_ − *r*_1_ − *a*_2_ − *r*_2_ system. Likewise, the addition of the effective diffusion terms does not significantly change the learned matrix. This indicates robustness of the reconstruction procedure (SI Appendix). The phase portraits of various learned sub-systems are shown in Fig. S1.

3) The idea of the reconstruction algorithm that we use is similar to the SINDy method [57, 58]. For our data, SINDy generated inconclusive results (SI Appendix), most likely because our system is close to a single stable steady state and dominated by noise.

4) The simplest type of dynamical system that we are reconstructing corresponds to Markovian dynamics without memory. In principle, because there are likely hidden biological variables unaccounted for in the model (see Discussion), the data could be better explained by delay differential equations, in which the change of the system state depends on its history, not just on the current state. However, a more advanced reconstruction with delay terms led to non-robust and non-interpretable results (SI Appendix).

4) We compared the reconstructed system based on the whole available data with the systems obtained from separate analyses of individual cells, and found that cell-to-cell variation of the reconstructed coefficients is not great and, on average, agrees with the whole-data approximation (Fig. S4, SI Appendix).

### The reconstructed linear stochastic dynamics predict experimentally measured fluctuations and correlations

To test that the reduced linear model reproduces the statistical features of the data, we simulated the reconstructed system with the learned noise and computed probability distributions of F-actin, Arp2/3, and velocity. Fig. 2A shows that the experimentally measured distributions (blue) are close to those from the simulations (orange). All 1D marginal distributions, including those for *v, a*_1_, *a*_2_, *r*_1_, and *r*_2_, exhibit unimodal, symmetric shapes that closely resemble Gaussian distributions (Fig. 2A, Fig. S5). This supports the modeling assumption that these variables follow linear stochastic dynamics subject to additive white noise. Such behavior is consistent with an Ornstein-Uhlenbeck process, the prototypical linear stochastic differential equation that produces stationary Gaussian fluctuations.

The 2D joint distributions (scatter plots with overlaid marginal densities) between F-actin and velocity, F-actin and Arp2/3, and Arp2/3 across layers reveal elliptical shapes, indicating strong pairwise correlations (Fig. 2A, Fig. S5). The orientation and elongation of these ellipses indicate pairwise correlations that are the same in the data and simulations, further validating the inferred dynamics.

To further probe these correlations, we computed ensemble-averaged cross-correlation functions and observed that the model semi-quantitatively reproduces experimentally measured cross-correlations (Fig. 1B, Fig. S6). For example, Fig. 2B (left) shows a small negative peak at a negative time lag, indicating that downward fluctuations in actin concentration (*a*_1_) precede changes in edge velocity, in agreement with the predicted negative feedback from actin density to the protrusive activity. The more prominent positive peak of the crosscorelation function with the positive time lag indicates the predicted strong positive feedback from the velocity to F-actin. Similarly, Fig. 2B (right) shows that Arp2/3 at the rear positively correlates, with a lag, with Arp2/3 at the front, consistent with the retrograde flow transporting Arp2/3 to the rear.

Crosscorrelations shown in Fig. S6 offer additional insights: 1) velocity almost immediately correlates with Arp2/3 at the leading edge; 2) F-actin almost immediately locally correlates with Arp2/3; 3) F-actin fluctuations near the leading edge precede those at the rear, hinting that retrograde flow transports actin from the front to the rear.

Together, these results confirm that the measured fluctuations of actin, Arp2/3, and edge velocity are consistent with the model-generated dynamics. By themselves, these distributions and correlations offer limited mechanistic insight, so at this point, the linear reconstructed dynamics have to be analyzed by more heuristic and less formal methods.

### Building nonlinear spatial-temporal model for Arp2/3 and F-actin densities

It is a trivial mathematical fact that different nonlinear dynamical systems may have the same steady states, and even the same linear approximations in the vicinity of these states. Thus, there is no unique way to determine which feedbacks maintain the lamellipodial dynamic equilibrium. One additional challenge is that we want to build the spatial-temporal model of the lamellipodium, while so far we have a compartmentalized model – essentially, the crudely discretized system of some partial differential equations. The following non-rigorous arguments based on parsimony guide us to the full nonlinear model.

Let us start with the equation for F-actin density. Linear reconstructed equations for F-actin in the leading and trailing compartments have the form (Fig. 2C):

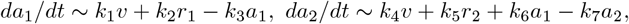

where *k*_*i*_, *i* = 1 − 7 are positive coefficients. Terms proportional to *r*_1_, *r*_2_ are easy to explain: these are source terms for actin filaments growing from Arp2/3 complexes. Terms proportional to *a*_1_, *a*_2_ could be responsible for F-actin disassembly. But what about the terms proportional to velocity? In principle, there could be subtle nonlinear effects according to which the actin density at the edge is an increasing function of velocity [40, 62]. However, we prefer to make as few such assumptions, not directly supported by our data, as possible. And then, how do we explain the effect of *v* on F-actin at the rear, *a*_2_?

Turns out that the process of retrograde flow of the actin network gives a natural answer to all these questions: Capped actin filaments are carried away from the edge with velocity *V*_*p*_ = (*V* + *V*_*f*_ ) in the framework fixed at the edge. Indeed, in the lab coordinate system, the edge moves forward with velocity *V*, and the actin network flows retrogradely with velocity *V*_*f*_, so relative to the edge the network flows retrogradely with velocity *V*_*p*_ = (*V* + *V*_*f*_ ), where *V*_*p*_ is the rate of actin filaments polymerization at the edge. In the model, we assume that the retrograde flow velocity *V*_*f*_, which we do not measure, is constant, and the protrusion velocity *V*, which is measured, is variable.

In the compartmentalized lamellipodium, it takes time ∼ *L/*(*V* + *V*_*f*_ ) for filaments to flow from leading to trailing compartment (*L* is the compartment size), so in the equation for F-actin density in the leading compartment, *A*_1_, there will be the term *dA*_1_*/dt* ∼ −((*V* + *V*_*f*_ )*/L*)*A*_1_. Note that even such simple term is nonlinear, as both *A*_1_ and *V* are dynamic variables. If we denote the stable steady states for actin and velocity as 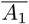 and 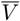 and small deviations from the steady states as *a*_1_ and *v*, respectively 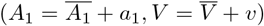, then the linearized equation for the leading compartment actin will have the respective terms: 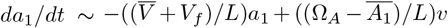, where Ω_*A*_ is the F-actin density at the very leading edge. This can explain the term proportional to the velocity in the reconstructed linear equation for the leading compartment actin density.

Similar arguments for the trailing compartment actin density give the following terms in respective linearized equation: 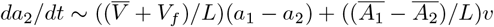 Not only does this equation explain the positive term proportional to the protrusion velocity (retrograde flow brings more actin from the front when protrusion is faster), but also the positive term proportional to the actin density in the leading compartment (retrograde flow brings more actin from the front when there is more actin at the front).

Altogether, this analysis suggests the following PDE for the actin density (see also Fig. 3C):

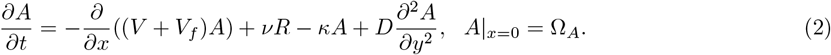

**FIG. 3.**
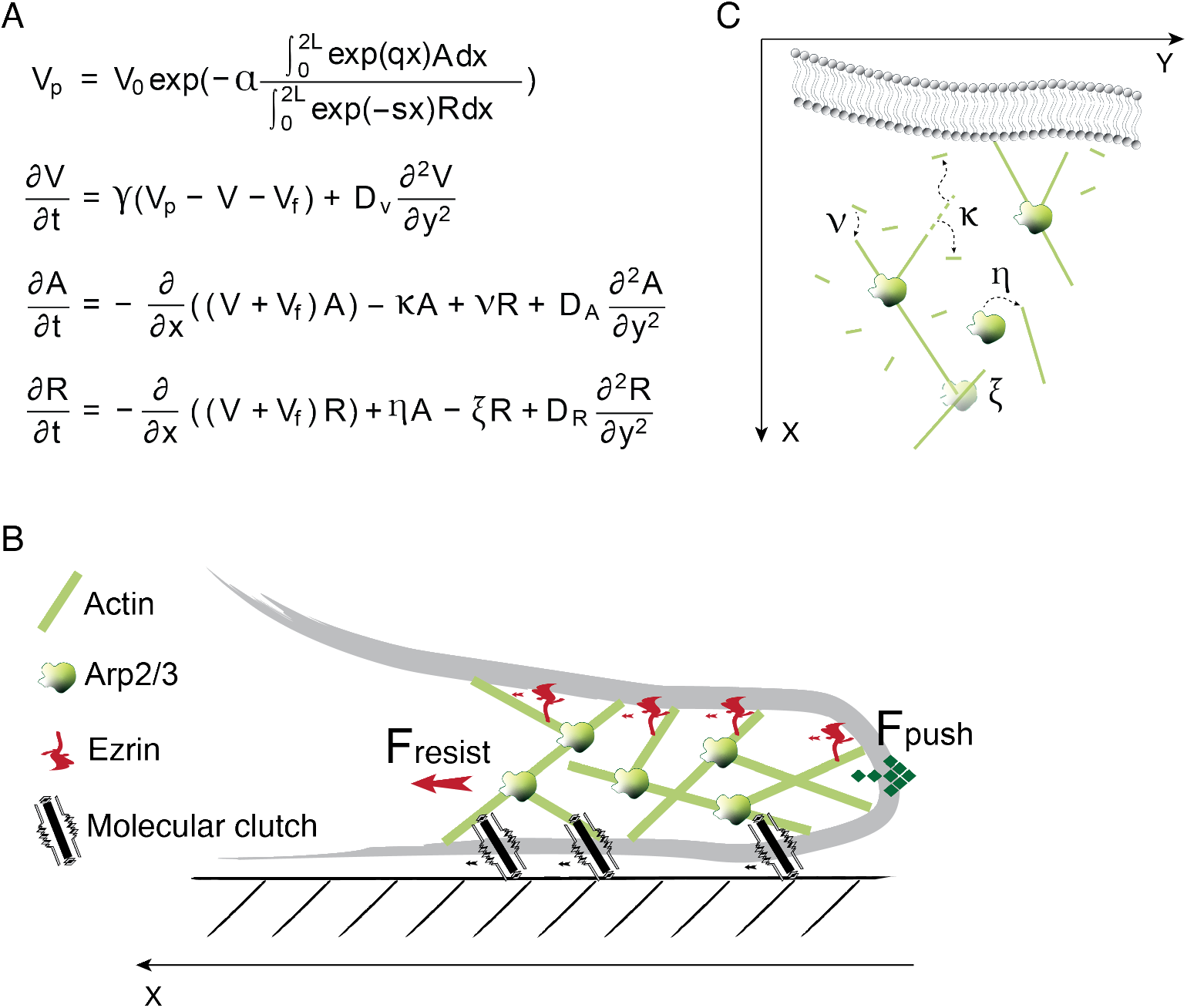
Nonlinear mechanistic model of protrusion dynamics. (A) Integro-PDE equations of the mechanistic model. Explanations are in the text. (B) Schematic illustration of the lamellipodial mechanics (view from the side). Growing actin filaments exert pushing forces on the membrane forward locally, at the leading edge. Ezrin and other actin-binding proteins transiently link actin filaments to the membrane, and molecular clutches of adhesion complexes link F-actin to the substrate, generating a global resistance force. (C) Schematic illustration of four linear reaction terms in the lamellipodia (view from the top). Actin filaments produce pushing forces at the leading edge. F-actin assembly on Arp2/3 rate is *ν*; its disassembly rate is *κ*. Arp2/3 detaches with rate *ξ*; its binding rate to F-actin is *η*.

The explanation for the boundary condition is: a constant density of the actin filaments growing against the leading edge is proportional to the constant Arp2/3 density (averaged over the narrow strip at the edge, see below) maintained by the cell at the edge.

Very similar arguments lead to the following PDE for Arp2/3 density:

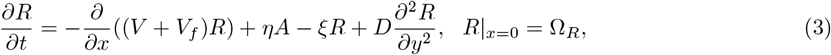

where Ω_*R*_ is a constant parameter equal to a constant Arp2/3 density at the leading edge that the cell maintains. What is the biophysical nature of the constant Arp2/3 density at the leading edge? We favor the following simple explanation: Arp2/3 complexes are activated at the very edge while being bound to WAVE molecules; when such activated Arp2/3 complex binds to the actin network, the complex starts moving away from WAVE with velocity *V*_*p*_. When the WAVE-Arp2/3 link stretches to a threshold, the link breaks, Arp2/3 is inserted into the network, and the next inactivated Arp2/3 complex binds to the freed WAVE molecule immediately and the process is repeated. In such a scenario, the Arp2/3 density at the edge is simply given by the WAVE density along the leading edge, and the rate of Arp2/3 insertion into the network is proportional to the retrograde flow rate.

Note that linearized reconstructed equations in Fig. 2C and full nonlinear Eqs. (2) and (3) indicate a positive feedback from protrusion velocity to both Arp2/3 and F-actin, which, we argue, is based on a very simple retrograde flow effect: the faster the protrusion is, the faster the high densities at the edge spread across the lamellipodium.

We discretized and scaled Eqs. (2) and (3), and found the model parameters from fitting the result to the reconstructed linear model (SI Appendix). This fitting is not unconstrained: from previously published data, we know the characteristic magnitude of parameter *V*_*f*_ [21], and from our data we know dimensional values for 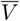 and *L*. Moreover, from our data we know the characteristic ratios of 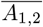 and 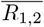 to the average values of *a*_1,2_ and *r*_1,2_ (SI Appendix). We found that the estimated coefficients obey all physical constraints from the data (SI Appendix), which is a testament of the model consistency.

The main conclusion of the model so far is that the only necessary (and sufficient) nonlinear terms in the molecular density equations are the drift terms. Respective retrograde treadmill is firmly established in the literature [21, 22, 27]. Estimates of the reaction rates (SI Appendix) offer several additional insights:

1. Actin disassembly rate, *κ*, is on the order of 1/min and is roughly uniform across the lamellipodium. Considering that the estimated time of the treadmill, 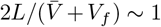 min, actin disassembles on the same scale as it drifts to the rear. According to measurements in [21], more than 80 % of the actin network is disassembled in the lamellipodium during the time of its treadmill to the rear, which agrees with the model estimate.
2. There is a significant branching action in the lamellipodium with rate *ν* ∼ 1/min meaning that a nascent filament is branched off from an Arp2/3 complex away from the edge within a minute. This could mean either that some Arp2/3 complexes are activated and associate with the actin network away from the edge (though the Arp2/3 majority attach to F-actin at the edge), or that rebranching (sequential filament disassembly on an Arp2/3 and then reassembly of another filament on the same Arp2/3) takes place across the lamellipodium [19]. Notably, rate *ν* is twice higher in the leading half of the lamellipodium.
3. Arp2/3 barely dissociates in the leading half of the lamellipodium, while in the trailing half, respective dissociation rate jumps to *ξ* ∼ 1.5/min.

Rate *η* of the actin-dependent Arp2/3 binding away from the edge was predicted to have an unusual behavior according to the linear reconstructed dynamics: negative closer to the edge (so higher F-actin would inhibit Arp2/3 binding) and positive closer to the rear (making F-actin a partially limiting substrate). However, in the best fit to the full model, parameter *η* is almost zero closer to the edge, so the hard to interpret inhibitory F-actin effect is absent from the nonlinear model. Closer to the rear, parameter *η* is small, ∼ 0.5/min. Indeed, F-actin is unlikely to be the rate-limiting factor for branching at the leading edge according to indirect data in [24].

### Building a nonlinear spatial-temporal model for the edge velocity

The reconstructed linear ODE for the protrusion velocity (Fig. 2C) indicates that the velocity is an increasing function of Arp2/3 and a decreasing function of the F-actin density, especially of the F-actin density at the lamellipodial rear. The most natural interpretation of this result is based on the well-established force-limited nature of the actin polymerization velocity at the lamellipodial leading edge [11, 23]. Namely, the simplest expression for the actin polymerization rate is the force-velocity relation: *V*_*p*_ = *V*_0_ exp(−*f/f*_0_), where *V*_0_ is the free polymerization rate determined by biochemical factors, multiplied by the exponential mechanical factor slowing down the polymerization, where *f* is the load force per filament and *f*_0_ is a constant force parameter defined by many structural and mechanical factors. The load force per filament is equal to the resistance force per unit length of the leading edge divided by the number of pushing filaments per unit length of the edge. The former, according to the learned linear model, is proportional to the integral of the actin density from the front to the rear, with the rear contributing more, so we chose to model the actin contribution to the resistance as follows: 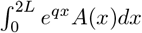. Here the exponential factor is responsible for the growing increase to the resistance of the F-actin density with distance from the edge.

The number of pushing filaments per unit length of the edge is most likely proportional to the number of Arp2/3 complexes near the edge, where the filaments branched off by these complexes reach the edge uncapped and push it forward (Fig. 3B). We model respective factor by the integral 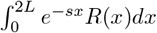 where *s*^−1^ is the average actin filament’s length on the order of a few hundred nanometers. Thus, we posit the following expression:

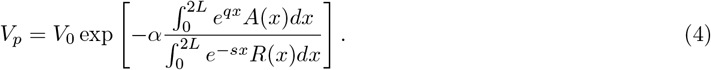

Here *α* is a non-dimensional (because we measure the Arp2/3 and F-actin densities in the same units proportional to numbers of complexes and filaments, respectively, per square micron) parameter.

The reconstructed linearized equation for velocity has a large negative coefficient in front of the velocity term indicating that the velocity rapidly relaxes to the quasi-steady state level defined by the local Arp2/3 density near the edge and global actin density, *V* → *V*_*p*_ − *V*_*f*_ . Thus, we posit the following dynamic equation for the protrusive velocity:

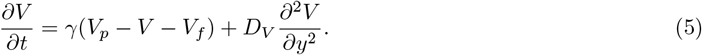

Here *γ* ∼ 0.08 sec^−1^ is respective relaxation rate. Considering the observed ∼ 15 nm/sec magnitude of the protrusion velocity fluctuations, this relaxation rate is consistent with viscoelastic cytoskeletal relaxation [63] on the microscopic scale of a single actin filament (∼ 15nm/sec ×*γ*^−1^ ≈ 200 nm is the characteristic average filament length [4]).

We discretized and scaled Eqs. (4) and (5) and found the model parameters from fitting the result to the reconstructed linear model (SI Appendix). Several insights emerged from the parameter estimates:

The free polymerization velocity is slowed down by an order of magnitude at the edge (Table. S1) indicating that an average resistive load force of a few pN per filament is applied on the leading edge, which is in agreement with experiment [23].

1. There is a simple explanation for why the resistance force is global and proportional to the actin density: actin filaments flowing back from the edge transiently attach through a host of actin binding proteins to both ventral and dorsal plasma membrane [62] and to adhesion complexes [22, 26–28, 30] (Fig. 3B) generating integrated drag slowing down the protrusion. Notably, the adhesion distribution is graded with the distance from the leading edge [22, 28], which can explain the distance dependence of the effective resistance. Note that this explanation implicitly assumes a relatively rigid lamellipodial actin network flowing retrogradely with a constant rate.
2. The model fit to the data allows estimate of the sum of the mean protrusion velocity 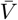 and retrograde flow rate *V*_*f*_ but not of these parameters separately. When we estimated this sum and subtracted the approximate experimentally measured dimensional parameter 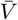 from the sum, the difference *V*_*f*_ was on the order of magnitude established in the literature. In the future, accurate measurements of the actin flow together with the protrusion velocity and protein concentrations will be helpful for further testing and refining the model.
3. Lastly, note that the posited equation for the protrusion velocity is not only highly nonlinear, but also nonlocal due to the global effect of F-actin on the mechanical resistance. This makes the full model not just the PDE model, but the integro-PDE model.

### Learned dynamics exhibits damped oscillations

Phase portraits of the reconstructed lamellipodial behavior (Fig. 1C, Fig. S1) hint at damped fluctuations near the dynamic equilibrium. The full spatial-temporal dynamics also exhibits oscillations (Fig. 4). To demonstrate the predictive capabilities of the mechanistic model, we numerically solved a full deterministic (without the noise) PDE system (Fig. 3A) with several nontrivial initial conditions. We found that the Arp2/3 and F-actin densities relax to the stable steady state, in which the densities are constant along the edge and decrease away from it. Then, we did a numerical “optogenetic” experiment, making Arp2/3 density equal to zero on a square domain starting at the edge and extending to the half-lamellipodial width and using such a state as the initial condition. Numerical solution illustrates how the Arp2/3-depleted region is transported back by the retrograde flow, transiently damping F-actin locally (Fig. 4). The lateral diffusion, meanwhile, slowly transports actin and Arp2/3 into the perturbed drifting domain, while Arp2/3 recovers at the edge. The computed edge velocity at the center of the perturbed region briefly decreases because, without initial Arp2/3 at the edge, there are too few filaments pushing the edge forward. As Arp2/3 density at the edge is rebuilt, the velocity not only recovers but overshoots upward because the collapsed overall actin density at the center exerts lower resistance. Then, the actin density recovers, and after small transient oscillations, the equilibrium is reached.

**FIG. 4.**
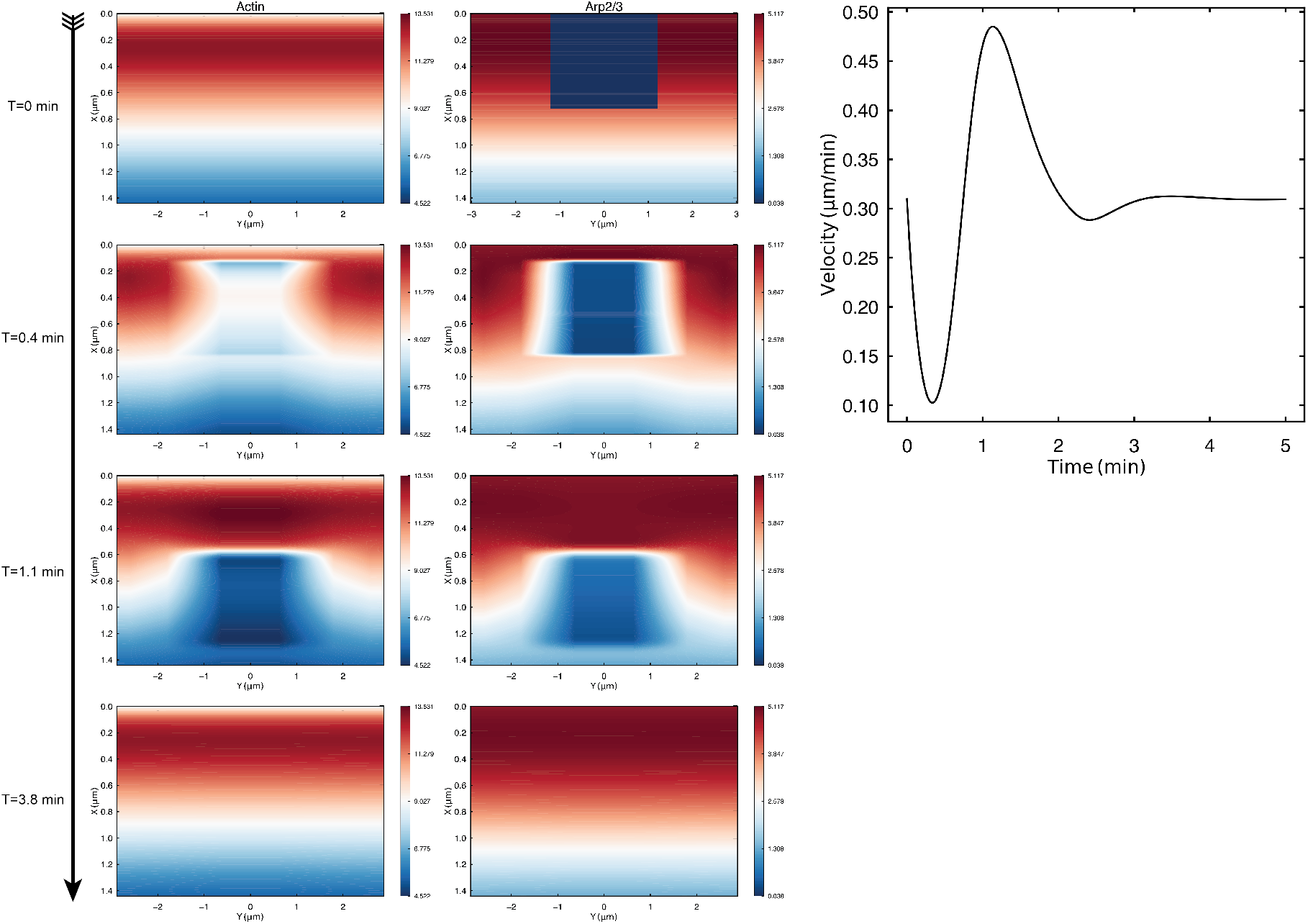
Simulated spatiotemporal dynamics of actin, Arp2/3, and velocity exhibits damped waves and oscillations. Left: Numerical solution of the integro-PDE model shown in Fig. 3A. From top to bottom, four snapshots of the spatial distribution of F-actin (left column) and Arp2/3 (right column). The color-coded densities are measured in non-dimensional units (SI appendix). Initial conditions correspond to an imaginary ‘optogenetic’ experiment: F-actin is constant in the *y*-direction (parallel to the leading edge) and linearly decreases to the rear; Arp2/3 is the same, but in addition is depleted in one of the leading compartments. Boundary conditions and other details are described in the text. Right: Computed (in the simulation shown at the left) protrusion velocity (as function of time) at the center of the leading compartment with initial damped Arp2/3 density.

We also calculated the spectrum of the reconstructed 5-dimensional *V* − *A*_1_ − *R*_1_ − *A*_2_ − *R*_2_ linear dynamical system characterizing a pair of leading and trailing compartments. All eigenvalues had negative real parts, but several also had imaginary parts. To illustrate the mechanical cycle in this pair of the leading and trailing compartments, we plotted the time evolution of *V* −*A*_1_ −*R*_1_ −*A*_2_ −*R*_2_ variables computed from the full nonlinear model, discretized and limited to two compartments, without diffusion and stochastic terms (Fig. 5A). We chose the initial conditions to be one of the eigenvectors of the linear dynamic coefficients matrix. Dominating relaxation terms then drive all of them back to the equilibrium, but subtle fluctuations emerge: F-actin at the leading edge inches upward briefly because of the positive feedbacks from the velocity and Arp2/3; high initial velocity transports Arp2/3 and F-actin to the rear generating transient increases of Arp2/3 and F-actin at the rear; the velocity overshoots downward before the equilibrium is reached because this transient increase of F-actin at the rear elevates the mechanical resistance.

**FIG. 5.**
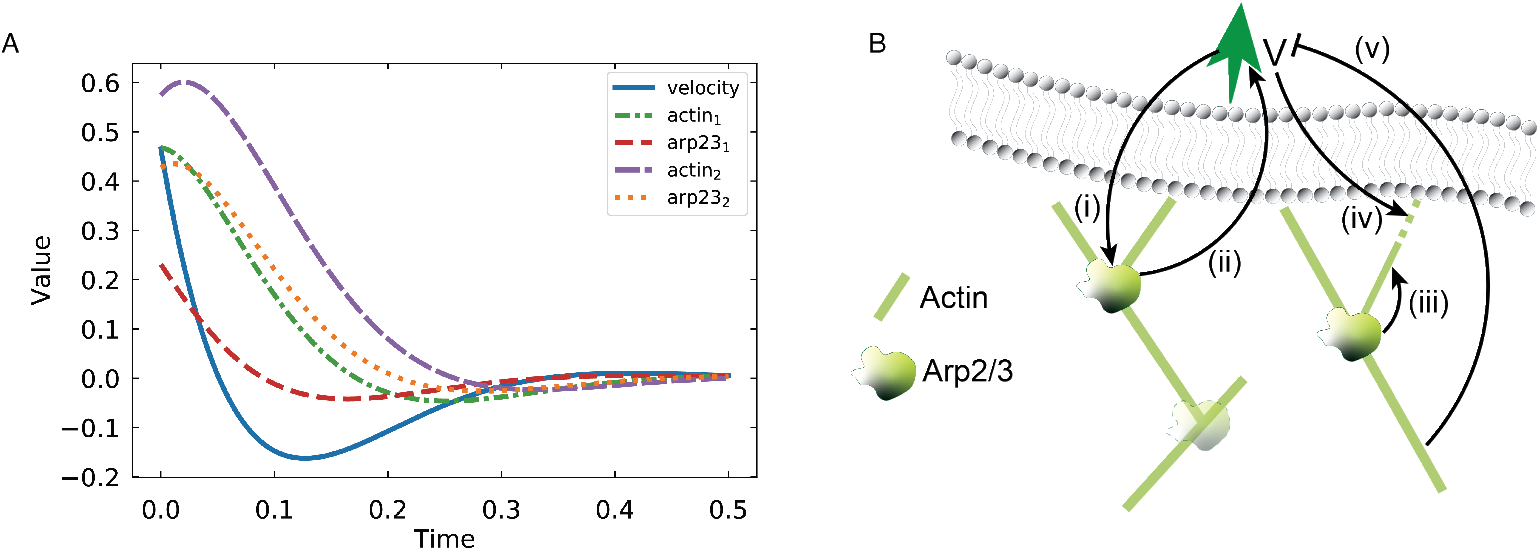
Principal feedbacks of the protrusion dynamics. (A) Simulated relaxation of coupled velocity, Arp2/3, and F-actin densities in a pair of leading and trailing compartments. The solution is found by numerically solving the full nonlinear model with initial conditions given by the real part of the eigenfunction of the linear system corresponding to the largest (by magnitude eigenvalue). The model was discretized and limited to the pair of leading and trailing compartments, thereby neglecting lateral diffusion. Stochastic terms were not simulated. (B) Principal feedbacks of the protrusion dynamics: (i) increased velocity increases the rate of Arp2/3 attachment ; (ii) Arp2/3-generated filaments push the edge forward; (iii) more Arp2/3 branches off more filaments; (iv) higher velocity accelerates F-actin growth; (v) higher F-actin density elevates mechanical resistance, slowing the protrusion down.

## DISCUSSION

One of the main results of this study is estimating of the dimensional values for principal reactions and mechanical and transport processes in the lamellipodium (Table. S1) from a single non-perturbing experiment. The learned mechanistic model is intuitive and conceptually simple, suggesting that the only essential nonlinearities in the lamellipodial dynamics stem from the unambiguous drift terms proportional to the velocity-density products. The main broad prediction of the model is that the lamellipodial dynamics is dominated by the following mechanical cycle: Arp2/3 complexes at the edge produce filaments pushing the edge forward; faster leading edge protrusion leads to faster treadmill and spreading of higher F-actin density across the whole lamellipodium, This elevates the mechanical resistance and slows down the protrusion (Fig. 5B). This cycle leads to damped mechanical oscillations.

Granger Causality approach predicted similar feedbacks [51] based on the same data. The only difference is that the analysis in [51] posited no statistically significant feedback between Arp2/3 to velocity. One possible explanation is that our mechanistic model suggests the incoherent feedforward loop from Arp2/3 to velocity (Arp2/3 activates velocity, but also activates actin, which inhibits velocity), so these two conflicting parallel pathways can partially cancel each other. The feedback from velocity to Arp2/3 could be masked by the combination of activating and inhibiting effects discussed above. This underscores the benefit of using both mechanistic and causality approaches to increase confidence in biological knowledge, as both approaches share the same limitation of not always resolving competing interactions in systems with biological redundancy [50, 64, 65].

The model suggests that: 1) Actin growth velocity is driven by uncapped growing actin filaments at the edge, the number of which is proportional to the number of Arp2/3 complexes at the front. Filament growth is proportional to protrusion velocity, in agreement with [66].

2) Actin assembly and Arp2/3 attachment occur largely near the edge; F-actin and Arp2/3 then drift to the rear with retrograde flow. Actin disassembly and Arp2/3 detachment take place on the drift time scale. Actin disassembly rate is almost uniform across the lamellipodium, while Arp2/3 detachment occurs mostly at the lamellipodial rear. These conclusions agree with [9, 67].

3) According to the model, mechanical resistance to the protrusion is proportional to the global F-actin density, with higher effective resistance from the rear. We posit that this resistance stems from transient, F-actin-limited (in agreement with [62, 68, 69]), attachments to the membrane and substrate.

### Nature of the noise in the lamellipodium

Experimentally measured time series of Arp2/3 concentration in one of the leading edge compartments reveal large-amplitude fluctuations (Fig. S7B). According to the model and data, the measured noise (standard deviation/mean) for the Arp2/3 signal from one lamellipodial window is in the tens of percent range. Given that the average number of Arp2/3 molecules per 0.7*µ*m× 0.7*µ*m-area compartment is on the order of hundreds [70], simple Poisson statistics would predict a noise lower than 10%, suggesting that Arp2/3 noise cannot be fully explained by Poisson statistics alone (Fig. S7B).

In the SI Appendix, we explore a fully stochastic model based on the full reconstructed model in this study, in which a small number of WAVE molecules at the leading edge turn over slowly and activate Arp2/3 complexes which, when activated, attach to the actin network and treadmill to the rear with it. Simulations of this model (SI Appendix) show that ∼ 15 WAVE molecules per micron of the edge turning over on the scale of minutes would explain the observed level of noise of Arp2/3 numbers: slow Poisson noise (with standard deviation 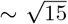 and mean ∼ 15) in the number of WAVE molecules per micron would propagate into similar relative fluctuations of the number of Arp2/3 molecules and then into respective fluctuations of the number of actin filaments and velocity.

The characteristic distance between nucleation promoting factors was measured to be about 25 nm [71], which gives about 40 WAVE molecules per micron of the edge, if they are placed in one row along the edge, which is not far from our estimate. In another study, however, hundreds of WAVE molecules were observed in multiple rows along a micron of the leading edge [10], with respective turnover rates reported in the range of seconds [9, 10], which could mean that fluctuations of signaling molecules’ concentrations upstream of WAVE is the main cause of the high lamellipodial noise.

### Effective lateral diffusion in the lamellipodium

An important insight from the model reconstruction is the learned effective diffusion of the protein densities and velocity along the leading edge. This effect is crucial for the stability of the edge – keeping the edge straight. Without the effective diffusion, protrusion/retraction fluctuations could make the edge uneven, splitting the smooth global protrusion into many transient uncoordinated local ones. Notably, ablation analysis in [51] showed that the lateral connections between variables in neighboring lamellipodial windows are the single most important factor for robust dynamic behavior. Recent study [45] suggests that the lamellipodial stability is underlined by the lateral actin flow [45, 72]: when actin filaments pushing the edge at varying angles grow along the protruding edge, their plus ends slide sideways along the edge. The stochastic nature of branching and capping then generates effective diffusion of F-actin along the edge. However, this does not explain the effective lateral diffusion of Arp2/3 molecules that are part of the actin network.

The clue to the nature of effective diffusion comes from the fact that the learned diffusion coefficients for F-actin and Arp2/3 are very similar, suggesting that F-actin and Arp2/3 diffusion originates from the same random process. The simplest assumption is that random lateral shifts of the actin network, as it flows retrogradely [73, 74], generate the effective lateral diffusion of F-actin and Arp2/3. The following estimates suggest that the order of magnitude estimates of the effective lateral diffusion coefficients for Arp2/3 and F-actin are not too far from the learned value, ∼ 0.2 *µ*m^2^*/*min. If local lateral retrograde flow changes directions on the characteristic time *τ*_*l*_ ∼ *γ*^−1^ ∼ 0.3 min of the learned characteristic relaxation time of the actin network (Table. S1), and if the lateral flow rate is *V*_*l*_ ∼ 0.3*V*_*f*_ ∼ 0.2 *µ*m*/*min [73], then 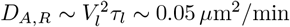 *µ*m^2^*/*min.

The most likely explanation for the effective diffusion of velocity is as follows: if velocity becomes higher at one point along the edge, then the local protrusion at that point pulls the membrane forward to the sides, relieving the load on pushing actin filaments and enabling easier and faster protrusions to the sides. Respective diffusion coefficient would be on the order of ∼ *V*_*p*_*l*_*a*_ ∼ 0.2 *µ*m^2^*/*min, where *l*_*a*_ ∼ 0.2 *µ*m is the characteristic length of an actin filament [2], which agrees with the order of magnitude of the learned parameter ∼ 0.55 *µ*m^2^*/*min.

These crude estimates are the best we can do: In the SI Appendix, we redid the regression analysis by coarsening the lamellipodial grid (lumping together several neighboring windows, Fig. S8) and found that the spatial resolution of the data does not allow learning more accurate values of the effective diffusion coefficients.

### Other possible interpretations of the learned model terms and caveats

It would be possible to attribute some terms in the learned model to complex nonlinear interactions of, for example, (i) Arp2/3 with actin [11, 18, 24], (ii) mechanochemical mechanism of accelerating actin assembly by stretching leading edge [66], (iii) autocatalytic velocity-Arp2/3-actin feedbacks [35, 40, 75]. Instead, we followed the principle of parsimony and ascribed nonlinearities to the drift process whenever possible.

The greatest uncertainty for interpreting mathematical terms is absence from the model of two prominent actin and velocity regulators at the leading edge, formin [51, 76, 77] and VASP [17, 51]. For example, our linear regression predicted negative feedback from F-actin to Arp2/3 at the front, which is in agreement with [78], where this effect was reported to be likely due to interactions with VASP. We plan on including formin and VASP fluctuation data [51] to Arp2/3-actin-velocity data in the next iteration of the model. Similarly, complex interactions between formin, F-actin and retrograde flow [79], or between Arp2/3-actin and adhesions or myosin [31, 80], or between actin polymerization and retrograde flow rate and adhesion mechanics [21, 74, 81], or regulated capping process [12] could be the cause of model terms that we excluded from the linear regression procedure. Also, Rho GTPases and other regulators upstream of Arp2/3, formin, and VASP are likely to be part of key pathways governing the leading edge dynamics [61, 82, 83], and their inclusion in future iterations of the model will be a must.

The negative feedback from F-actin to velocity could, in principle, be due to a limited amount of G-actin and conservation of the total actin amount in the lamellipodium: more F-actin leads to less G-actin and slower protrusion [33]. However, this is unlikely because a large G-actin concentration not limiting the protrusion was reported [70]. Nevertheless, nontrivial modes of G-actin and Arp2/3 delivery to the cell edge [14, 71] could influence interpretations of the model terms.

Less conceptual, more technical limitation of the model stems from neglecting variable dynamic curvature of the leading edge [51]. Subtle effects of coupling of leading-edge geometry and F-actin density [45] could introduce quantitative corrections to the learned mathematical terms. Another unaccounted-for factor is the heterogeneity of the retrograde flow of the deformable actin network [74]. These factors are unlikely to qualitatively change the model.

## Data, Materials, and Software Availability

All study data are included in the article and/or SI Appendix.

## Acknowledgements

We thank A. Manhart and K. Keren for useful suggestions. W.S. and A.M. were supported by NSF grant DMS 1953430. C.E.M. was supported by NSF CAREER DMS-2545859 and NSF/NIGMS DMS-2451263. J.N. and G.D. were supported by NIH grants R35GM136428.

## METHODS

### Experimental Data

We used the data on the spatiotemporal dynamics of the lamellipodial leading edge reported in [51], where U2OS cells co-expressing HaloTag-labeled Arp2/3 and mNeonGreen-labeled actin were analyzed. There, a window-based tracking system that segments the lamellipodium into a grid was employed, and time series for local edge velocity (*V* ), F-actin density (*A*), and Arp2/3 density (*R*) at 3-second intervals were extracted. The assumption is that the fluctuations reflect only F-actin density and Arp2/3 molecules in the network, while actin monomers and Arp2/3 in cytoplasm contribute to the uniform background signal only. Low-frequency fluctuations were filtered away from these series. The data were then normalized independently for each cell, such that each variable has a zero mean and unit variance. Further details are in the SI Appendix.

### Data-Driven Linear Reconstruction

We first reconstructed the system dynamics using a data-driven linear stochastic partial differential equation (SPDE) model to identify coupling terms unbiasedly. The state vector *u*(*y, t*) = (*v, a*_1_, *r*_1_, *a*_2_, *r*_2_)^*T*^ represents the normalized fluctuations of velocity and protein densities across two spatial layers. The resulting SPDEs can be described as 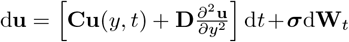. The evolution of the system is governed by a linear interaction matrix **C**, a diagonal diffusion matrix **D**, and a diagonal matrix of white noise strength ***σ***. To determine the interaction coefficients, we employed LASSO (Least Absolute Shrinkage and Selection Operator) regression, which enforces sparsity to select only the most significant physical couplings. This reconstruction yielded a model containing 25 non-zero empirical coefficients that describe the effective linear dynamics of the system. Further details are in the SI Appendix.

### Mechanistic Continuum Model and Parameter Identification

To provide a biophysical interpretation of the learned dynamics, we utilized the nonlinear continuum model introduced in the Results. We then (i) scaled and nondimensionalized, (ii) discretized, and (iii) linearized the equations around their steady state. This derivation yields a Jacobian matrix where each element is a function of the physical parameters. To obtain the parameters of the nonlinear model, we then treated the problem as an optimization task, employing ordinary least squares to minimize the difference between the theoretical Jacobian elements and the empirical LASSO-derived coefficients. Further details are in the SI Appendix.

### Numerical implementation of the PDE Model

Results of Fig. 4 are generated by numerical solution of the integro-PDE equations presented in Fig. 3A with initial and boundary conditions described in the text. The spatial domain was discretized using Chebyshev nodes to facilitate high-accuracy evaluation of the integral term via Clenshaw-Curtis quadrature. For the spatial differential operators, we used a five-point finite difference stencil adapted to the non-uniform Chebyshev grid, with standard finite difference formulas used to approximate the drift and diffusion terms. Temporal integration was performed using the backward Euler method. We verified the solution’s convergence by mesh refinement.

### Stochastic Simulation of Molecular Noise

We implemented a stochastic version of the mechanistic model that is described in the appendix.

## Notes

### Competing Interest Statement

The authors have declared no competing interest.

