## Supplemental materials for "Reconstructing Actin Dynamics of the Leading Edge from Observational Data"

### Supplementary Information
### for
### Reconstructing Actin Dynamics of the Leading Edge from Observational Data

Wenzheng Shi,[1] Christopher E. Miles,[2] Jungsik Noh,[3] Gaudenz Danuser,[4] and Alex Mogilner[5,*]

[1]Courant Institute, New York University, New York, NY 10012, USA

[2]Department of Mathematics, University of Utah, Salt Lake City, UT 84112, USA

[3]Lyda Hill Department of Bioinformatics, UT Southwestern Medical Center, Dallas, TX 75390, USA

[4]Lyda Hill Department of Bioinformatics, UT Southwestern Medical Center, Dallas, TX 75390, USA
Institute of Human Biology, Basel, Switzerland

[5]Courant Institute and Department of Biology,
New York University, New York, NY 10012, USA

(Dated: January 31, 2026)

#### DATA AND METHODS

##### Time series of protrusion velocity, F-actin, and Arp2/3

The data used in this paper were reported in [1]. The data were acquired from fluorescence time-lapse images of U2OS cells co-expressing HaloTag-labeled Arp2/3 and mNeonGreen-tagged actin. Imaging was performed at 3-second intervals over 15 minutes. The imaged lamellipodia were segmented into a grid of approximately square windows (compartments) that track with the moving edge. Given an average lamellipodial depth of 1.4 $\mu$m, a computational domain extending 1.4 $\mu$m from the edge, divided into two layers of $720 \times 720$ nm$^2$ windows, was defined. Time series of Arp2/3 and actin fluorescence intensities were extracted from each window. Simultaneously, local edge velocities were calculated from the displacement of the front-most windows, with positive and negative values corresponding to protrusion and retraction, respectively. As the raw intensity signals for F-actin and Arp2/3 exhibited low-frequency fluctuations attributed to global regulatory shifts in cells, a Fourier-domain high-pass filter was applied to isolate the relevant fast-scale dynamics. Specifically, the molecular time series were transformed into the frequency domain, where the first three lowest-frequency modes were zeroed out before inverse transformation.

##### Reconstructing a system of linear SPDEs from multi-component time series signals

To infer a dynamical model describing the interactions between edge velocity, F-actin, and Arp2/3, we utilized the space-time-resolved time series extracted from fluorescence imaging. Because these signals vary in physical units (velocity in nm/s; F-actin, and Arp2/3 – in arbitrary fluorescence units), we first normalized the data independently for each cell; specifically, the ensemble mean and standard deviation across the spatial com-

partments were computed and used to rescale the variables, so that the time series have zero mean and unit variance.

We began by using a system of coupled linear ordinary differential equations (ODEs). Specifically, we modeled the local rate of change of each variable as a linear combination of all measured variables in the same windows. We denoted the variables as follows: $v$ for velocity, $a_1$ and $a_2$ for F-actin in the leading and trailing window, and $r_1$ and $r_2$ for Arp2/3 in the leading and trailing window. The linear ODE system is written as:

$$\frac{\mathrm{d}}{\mathrm{d}t}\mathbf{u}(y,t) = \mathbf{C}\mathbf{u}(y,t), \tag{S1}$$

where $\mathbf{u} = (v, a_1, r_1, a_2, r_2)^T$ and $\mathbf{C}$ is a $5 \times 5$ matrix of unknown coefficients $c_{ij}$.
To estimate the coefficients, we discretized time using forward differences:

$$\frac{v^{t+1} - v^t}{\Delta t} = c_1 v^t + c_2 a_1^t + c_3 r_1^t + c_4 a_2^t + c_5 r_2^t, \tag{S2}$$

and analogous equations for the other variables. The time step $\Delta t = 0.01$: we use the characteristic time scale of 300 sec (the characteristic time of the network drift across the lamellipodium is on the order of minutes) as the unit of time, so the dimensional value $\Delta t = 3$ sec corresponds to $\Delta t = 0.01$ in the chosen time units. Five terms in the ODEs that are hard to interpret biologically (see the main text) are set to zero, i.e., $c_5 = 0$ in Eq. (S2).
We next extended the model to account for spatial interactions by introducing lateral diffusion terms, yielding a system of linear partial differential equations (PDEs):

$$\frac{\partial}{\partial t}\mathbf{u}(y,t) = \mathbf{C}\mathbf{u}(y,t) + \mathbf{D}\frac{\partial^2 \mathbf{u}}{\partial y^2}, \tag{S3}$$

where $\mathbf{D}$ is a diagonal matrix of effective diffusivities $D_i$. Discretizing the diffusion term using central differences, the velocity equation becomes:

$$\frac{v^{t+1} - v^t}{\Delta t} = c_1 v^t + c_2 a_1^t + c_3 r_1^t + c_4 a_2^t + c_5 r_2^t + D_1 \frac{v_L^t - 2v^t + v_R^t}{\Delta y^2}, \tag{S4}$$

where $v_L^t$ and $v_R^t$ denote neighboring windows in the lateral direction. The window spacing $\Delta y$ is 720 nm.
Finally, we formulated a stochastic extension of the PDEs by adding an independent white noise term to each equation, resulting in a system of stochastic PDEs (SPDEs):

$$\mathrm{d}\mathbf{u} = \left[\mathbf{C}\mathbf{u}(y,t) + \mathbf{D}\frac{\partial^2 \mathbf{u}}{\partial y^2}\right]\mathrm{d}t + \boldsymbol{\sigma}\mathrm{d}\mathbf{W}_t, \tag{S5}$$

where $\boldsymbol{\sigma}$ is a diagonal matrix, denoting the noise amplitude for each component, and $\mathbf{W}_t$ represents standard Brownian motion. The resulting SPDEs provide a data-driven linearized model of lamellipodial dynamics. The full system with fitted coefficients is presented in Fig. 2C in the main text.

To determine the coefficients' values in this linear system, we employed LASSO (Least Absolute Shrinkage and Selection Operator) regression. This linear regression yielded a model that describes the effective linearized dynamics of the system near the stable steady state.

#### Constructing mechanistic model from the reconstructed linearized dynamics

Based on the SPDE system reconstructed from the data (Fig. 2C in the main text), we developed a mechanistic continuous model to describe nonlinear lamellipodial dynamics. This model aims to provide a coarse-grained biophysical interpretation of the observed spatiotemporal behaviors of protrusion velocity ($V(y,t)$), F-actin ($A(x,y,t)$), and Arp2/3 ($R(x,y,t)$) densities. For simplicity, we decouple stochastic and deterministic terms and here focus on the latter, so the model is a system of coupled PDEs:

$$\begin{aligned}
V_p &= V_0 \exp(-\alpha \frac{\int_0^{2L} \exp(qx) A dx}{\int_0^{2L} \exp(-sx) R dx}),\\
\frac{\partial V}{\partial t} &= \gamma(V_p - V - V_f) + D_V \frac{\partial^2 V}{\partial y^2},\\
\frac{\partial A}{\partial t} &= -\frac{\partial}{\partial x}((V + V_f)A) - \kappa A + \nu R + D_A \frac{\partial^2 A}{\partial y^2},\\
\frac{\partial R}{\partial t} &= -\frac{\partial}{\partial x}((V + V_f)R) + \eta A - \xi R + D_R \frac{\partial^2 R}{\partial y^2}.
\end{aligned} \tag{S6}$$

Here, $x$-axis is normal to the leading edge with $x = 0$ corresponds to the edge, and $x = 2L$ is the lamellipodial rear. $y$-axis is along the leading edge. $V_p$ is the variable F-actin polymerization velocity, $V_0$ is the constant free polymerization rate, $V_f$ is the constant retrograde flow velocity. Constant $\alpha$ is the non-dimensional mechanical resistance parameter (Arp2/3 and F-actin densities are measured in the same units, see below). Constant parameter $q$ is the inverse characteristic length on which the F-actin effect on the mechanical resistance increases toward the rear. Constant parameter $s$ is the inverse characteristic width of the zone near the edge where Arp2/3-branched actin filaments are uncapped and push the edge membrane; we assume that this width is on the order of an average actin filament length. Constant parameter $\gamma$ is the inverse velocity relaxation time. The advection terms represent retrograde flow of the actin network, including Arp2/3 molecules. Model parameters $\kappa$, $\nu$, $\eta$, and $\xi$ are the actin disassembly rate, Arp2/3-mediated actin assembly rate, actin-dependent Arp2/3 attachment rate and Arp2/3 detachment rate, respectively. These parameters, in fact, are spatially graded – they are functions of the distance from the leading edge, $\kappa(x), \nu(x), \eta(x), \xi(x)$. As the spatial resolution in the experiment is limited to just two compartments across the lamellipodial width, in Table S1 we provide two values for each of these parameters – one at the front, another at the rear of the lamellipodium. $D_V$, $D_A$, $D_R$ represent the effective lateral diffusivity of velocity, F-actin and Arp2/3, respectively.

We use no-flux boundary conditions at the lateral lamellipodial ends and Dirichlet boundary conditions for the F-actin and Arp2/3 densities at the lamellipodial edge:

$$\begin{aligned}
&R|_{x=0} = \Omega_R, A|_{x=0} = \Omega_A,\\
&\left.\frac{\partial V}{\partial y}\right|_{y=-\tilde{L},\tilde{L}} = \left.\frac{\partial A}{\partial y}\right|_{y=-\tilde{L},\tilde{L}} = \left.\frac{\partial R}{\partial y}\right|_{y=-\tilde{L},\tilde{L}} = 0.
\end{aligned} \tag{S7}$$

Here $2\tilde{L}$ is the lamellipodial length, which is much greater than the lamellipodial width $2L$. We discuss parameters $\Omega_R, \Omega_A$ below. Note that because of the constant given Arp2/3 and F-actin densities at the very edge, the fluxes of Arp2/3 and F-actin densities from the edge into the network are proportional to the velocity: $R|_{x=0} = \Omega_R \Rightarrow J_R|_{x=0} = \Omega_R(V + V_f)$, $A|_{x=0} = \Omega_A \Rightarrow J_A|_{x=0} = \Omega_A(V + V_f)$. Note also that the PDEs for variables $V, R, A$ are reaction-diffusion equations along $y$ direction, hence two boundary conditions at the left and right lamellipodial ends. The PDEs for $R, A$ are reaction-drift equations along $x$ direction, hence just one boundary condition at the edge from which the drift starts.

We nondimensionalized Eqs. (S6) and (S7) using characteristic length scale $L = 720$ nm (lamellipodium's half-width) and characteristic time scale $T = 300$ sec. For velocity, F-actin and Arp2/3 densities we used scales $\widetilde{V}, \widetilde{A}, \widetilde{R}$ equal to respective measured standard deviation (in units of nm/sec for velocity and fluorescent signal units for F-actin and Arp2/3 densities, see below). In principle, ratio $L/T$ could also be used as the velocity scale but this did not result in simplification. After nondimensionalization, Eq. (S6) become:

$$
\begin{aligned}
V_p &= \frac{V_0}{\widetilde{V}} \exp(-\alpha \frac{\widetilde{A}\int_0^2 \exp(qLx)Adx}{\widetilde{R}\int_0^2 \exp(-sLx)Rdx}) \\
\frac{\partial V}{\partial t} &= \gamma T(V_p - V - V_f) + \frac{D_V T}{L^2}\frac{\partial^2 V}{\partial y^2} \\
\frac{\partial A}{\partial t} &= -\frac{\widetilde{V}T}{L}\frac{\partial}{\partial x}((V+V_f)A) - \kappa T A + \frac{\widetilde{R}\nu T}{\widetilde{A}}R + \frac{D_A T}{L^2}\frac{\partial^2 A}{\partial y^2} \\
\frac{\partial R}{\partial t} &= -\frac{\widetilde{V}T}{L}\frac{\partial}{\partial x}((V+V_f)R) + \frac{\widetilde{A}\eta T}{\widetilde{R}}A - \xi T R + \frac{D_R T}{L^2}\frac{\partial^2 R}{\partial y^2}
\end{aligned}
\tag{S8}
$$

Here we kept the notations for the rescaled nondimensional variables $V$, $A$, and $R$ and for rescaled nondimensional parameters $V_f$, $\Omega_A$ and $\Omega_R$ unchanged.

Eq. (S8) were then discretized into two spatial layers across the width of the lamellipodium to mirror the experimental compartmentalization. Letting $A_1$, $A_2$, $R_1$, $R_2$ denote values in the leading and trailing windows, the equations become:

$$
\begin{aligned}
\frac{\partial V}{\partial t} &= \gamma T\left[\frac{V_0}{\widetilde{V}}\exp\left(-\alpha\frac{\widetilde{A}}{\widetilde{R}}\frac{s}{q}(\exp(qL)-1)\frac{A_1+\exp(qL)A_2}{R_1}\right) - V - V_f\right] + \frac{D_V T}{L^2}\frac{\partial^2 V}{\partial y^2} \\
\frac{\partial A_1}{\partial t} &= \frac{\widetilde{V}T}{L}(\Omega_A - A_1)(V+V_f) - \kappa_1 T A_1 + \frac{\widetilde{R}\nu_1 T}{\widetilde{A}}R_1 + \frac{D_A T}{L^2}\frac{\partial^2 A_1}{\partial y^2} \\
\frac{\partial A_2}{\partial t} &= \frac{\widetilde{V}T}{L}(A_1 - A_2)(V+V_f) - \kappa_2 T A_2 + \frac{\widetilde{R}\nu_2 T}{\widetilde{A}}R_2 + \frac{D_A T}{L^2}\frac{\partial^2 A_2}{\partial y^2} \\
\frac{\partial R_1}{\partial t} &= \frac{\widetilde{V}T}{L}(\Omega_R - R_1)(V+V_f) - \xi_1 T R_1 + \frac{\widetilde{A}\eta_1 T}{\widetilde{R}}A_1 + \frac{D_R T}{L^2}\frac{\partial^2 R_1}{\partial y^2} \\
\frac{\partial R_2}{\partial t} &= \frac{\widetilde{V}T}{L}(R_1 - R_2)(V+V_f) - \xi_2 T R_2 + \frac{\widetilde{A}\eta_2 T}{\widetilde{R}}A_2 + \frac{D_R T}{L^2}\frac{\partial^2 R_2}{\partial y^2}
\end{aligned}
\tag{S9}
$$

Assuming small perturbations around steady-state values, we linearized the system by substituting, for all dependent variables, $X = \overline{X} + x$, where $\overline{X}$ denotes the steady state and $x$ denotes linear perturbations. Substituting such expansions into Eq. (S9) and neglecting nonlinear terms gives the following linearized system:

$$
\begin{aligned}
\frac{\partial v}{\partial t} &= -\gamma T v + \gamma T(\overline{V}+V_f)\left(-\frac{\alpha\frac{\widetilde{A}}{\widetilde{R}}\frac{s}{q}(\exp(qL)-1)}{\overline{R_1}}\right)\left(a_1 + \exp(qL)a_2 + \left(-\frac{\overline{A_1}+\exp(qL)\overline{A_2}}{\overline{R_1}}\right)r_1\right) + \frac{D_V T}{L^2}\frac{\partial^2 v}{\partial y^2}, \\
\frac{\partial a_1}{\partial t} &= \frac{\widetilde{V}T}{L}(\Omega_A - \overline{A_1})v - \left(\frac{\widetilde{V}T}{L}(\overline{V}+V_f) + \kappa_1 T\right)a_1 + \frac{\widetilde{R}\nu_1 T}{\widetilde{A}}r_1 + \frac{D_A T}{L^2}\frac{\partial^2 a_1}{\partial y^2}, \\
\frac{\partial a_2}{\partial t} &= \frac{\widetilde{V}T}{L}(\overline{A_1}-\overline{A_2})v + \frac{\widetilde{V}T}{L}(\overline{V}+V_f)a_1 - \left(\frac{\widetilde{V}T}{L}(\overline{V}+V_f) + \kappa_2 T\right)a_2 + \frac{\widetilde{R}\nu_2 T}{\widetilde{A}}r_2 + \frac{D_A T}{L^2}\frac{\partial^2 a_2}{\partial y^2}, \\
\frac{\partial r_1}{\partial t} &= \frac{\widetilde{V}T}{L}(\Omega_R - \overline{R_1})v + \frac{\widetilde{A}\eta_1 T}{\widetilde{R}}a_1 - \left(\frac{\widetilde{V}T}{L}(\overline{V}+V_f) + \xi_1 T\right)r_1 + \frac{D_R T}{L^2}\frac{\partial^2 r_1}{\partial y^2}, \\
\frac{\partial r_2}{\partial t} &= \frac{\widetilde{V}T}{L}(\overline{R_1}-\overline{R_2})v + \frac{\widetilde{V}T}{L}(\overline{V}+V_f)r_1 + \frac{\widetilde{A}\eta_2 T}{\widetilde{R}}a_2 - \left(\frac{\widetilde{V}T}{L}(\overline{V}+V_f) + \xi_2 T\right)a_2 + \frac{D_R T}{L^2}\frac{\partial^2 r_2}{\partial y^2},
\end{aligned}
\tag{S10}
$$

where the steady states can be computed from solving these five algebraic equations for five steady state values:

$$
\begin{aligned}
\overline{V}+V_f &= \frac{V_0}{\widetilde{V}}\exp(-\alpha\frac{\widetilde{A}}{\widetilde{R}}\frac{s}{q}(\exp(qL)-1)\frac{\overline{A_1}+\exp(qL)\overline{A_2}}{\overline{R_1}}),\\
\overline{A_1} &= \frac{\frac{\widetilde{V}T}{L}\Omega_A\left(\overline{V}+V_f\right)+\frac{\widetilde{R}\nu_1 T}{\widetilde{A}}\overline{R_1}}{\frac{\widetilde{V}T}{L}(\overline{V}+V_f)+\kappa_1 T},\\
\overline{A_2} &= \frac{\frac{\widetilde{V}T}{L}\overline{A_1}\left(\overline{V}+V_f\right)+\frac{\widetilde{R}\nu_2 T}{\widetilde{A}}\overline{R_2}}{\frac{\widetilde{V}T}{L}(\overline{V}+V_f)+\kappa_2 T},\\
\overline{R_1} &= \frac{\frac{\widetilde{V}T}{L}\Omega_R\left(\overline{V}+V_f\right)+\frac{\widetilde{A}\eta_1 T}{\widetilde{R}}\overline{A_1}}{\frac{\widetilde{V}T}{L}(\overline{V}+V_f)+\xi_1 T},\\
\overline{R_2} &= \frac{\frac{\widetilde{V}T}{L}\overline{R_1}\left(\overline{V}+V_f\right)+\frac{\widetilde{A}\eta_2 T}{\widetilde{R}}\overline{A_2}}{\frac{\widetilde{V}T}{L}(\overline{V}+V_f)+\xi_2 T}.
\end{aligned}
\tag{S11}
$$

Note that, ideally, Eq. (S10) should have the same coefficients as those in the equations in Fig. 2C in the main text. However, the mechanistic model yields only 15 independent coefficients (without diffusion coefficients and noise amplitudes), while the SPDE system has 18 (without diffusion coefficients and noise amplitudes). To resolve this over-constrained system, we employed ordinary least squares to minimize the difference between two sets of coefficients, and the best-fit estimates are:

$$
\begin{aligned}
&\gamma T = 23.39, \quad \frac{V_0}{\widetilde{V}} = 24.29, \quad \alpha\frac{\widetilde{A}}{\widetilde{R}}\frac{s}{q}(\exp(qL)-1) = 0.53, \quad \exp(qL) = 3.45,\\
&\frac{\widetilde{V}T}{L} = 8.16, \quad \kappa_1 T = 6.24, \quad \kappa_2 T = 3.99, \quad \frac{\widetilde{R}\nu_1 T}{\widetilde{A}} = 9.40, \quad \frac{\widetilde{R}\nu_2 T}{\widetilde{A}} = 4.40,\\
&\xi_1 T = 0, \quad \xi_2 T = 7.76, \quad \frac{\widetilde{A}\eta_1 T}{\widetilde{R}} = 0, \quad \frac{\widetilde{A}\eta_2 T}{\widetilde{R}} = 2.81\\
&\Omega_A = 6.78, \quad \Omega_R = 3.49, \quad \frac{D_V T}{L^2} = 5.34, \quad \frac{D_A T}{L^2} = \frac{D_R T}{L^2} = 1.79,
\end{aligned}
\tag{S12}
$$

Respective dimensional parameters are shown in Table S1. Moreover, the steady states are found as follows:

$$
\overline{A_1} = 5.90, \quad \overline{A_2} = 4.64, \quad \overline{R_1} = 3.33, \quad \overline{R_2} = 2.41 \tag{S13}
$$

Several relevant notes:

1) The protrusion velocity appears in the model always in combination with the retrograde flow rate, as the sum $(\overline{V}+V_f)$. It is the evidence of the model validity that after this sum's value was found from fitting the reconstructed linear model, and parameter $V_f$ is chosen to be on the order of the published measurements: treadmilling rates of the retrograde flow are measured to be on the order of 10 nm per second [2], similar to the retrograde flow rates [3]; we chose parameter $V_f$ on the same order of magnitude in the model. The magnitude of the average protrusion velocity $\overline{V}$ turns out to be very similar to the measured dimensional value.

2) The velocity scale $\widetilde{V}$ was found from fitting the reconstructed linear model, but this parameter is also the measured dimensional standard deviation of the protrusion velocity (used in the process of reconstruction of the linear SODEs from the data). These two values turn out to be very close (Table S1), further attesting to the model consistency.

3) Arp2/3 and F-actin densities are measured in units of respective fluorescent signals in the experiment. It is not possible to convert these measurements accurately into physical units, but rough estimates are possible

along the following lines. First, we subtracted average background fluorescence (from the areas outside the cell) from the spatially varying lamellipodial fluorescence signal, in images similar to those in Fig. 1A of the main text, and interpreted the mean and standard deviation of this difference as the mean and standard deviation of Arp2/3 and F-actin densities, respectively. This was done with a small number of images using ImageJ software, and the result can only be used as a rough, order-of-magnitude estimate. We found that for both Arp2/3 and F-actin densities, the means are 3 to 5 times greater than the respective standard deviations. Note that these standard deviations are used as the scales of the respective densities, and so in all formulas above $\widetilde{A} = \widetilde{R} = 1$. The fact that $4 < \overline{A} < 6$ and $2 < \overline{R} < 4$ agrees with the rough experimental estimates of the mean to standard deviation ratio adds support to the model.

Second, there is one simple natural choice of the physical dimension which is the same for the Arp2/3 and F-actin densities: number (of Arp2/3 complexes and of actin filaments, respectively) per square micron. We prefer this choice, and so there are no ratios of actin to Arp2/3 densities in dimensional parameters $\nu$ and $\eta$.

Third, note that the model predicts a decrease of the Arp2/3 and F-actin densities from the front to the rear ($\overline{A}_1 > \overline{A}_2, \overline{R}_1 > \overline{R}_2$), which is in agreement with experimental observations [4].

4) The best fit to the reconstructed Arp2/3 detachment rate in the leading compartment was so small that in the Table S1 we list this rate simply as 0, as there would be no confidence in the exact value of this rate.

5) The best fit to the reconstructed coefficient $\eta_1$ in front of the actin-dependent inhibitory effect on Arp2/3 attachment in the leading compartment also turned out to be so small that in Table S1 we list this rate simply as 0, as there would be no confidence in the exact value of this rate. This is the only coefficient for which the approximation in the final full model is very poor.

TABLE S1. The dimensional parameters reconstructed through the mechanistic model. † value extracted from experiments, ‡ value computed from mechanistic model.

| Parameters | Interpretations | Values |
|---|---|---|
| $\gamma$ | velocity relaxation rate | $\approx 4.7$ min$^{-1}$ |
| $\widetilde{V}$ | standard deviation of velocity fluctuations | $^{\dagger} \approx 18$ nm $\cdot$ s$^{-1}$, $^{\ddagger} \approx 20$ nm $\cdot$ s$^{-1}$ |
| $V_0$ | free F-actin polymerization rate | $\approx 30$ $\mu$m $\cdot$ min$^{-1}$ |
| $\overline{V}$ | average edge protrusion velocity | $\approx 0.19$ $\mu$m $\cdot$ min$^{-1}$ |
| $V_f$ | actin network retrograde flow velocity | $\approx 0.70$ $\mu$m $\cdot$ min$^{-1}$ |
| $s$ | inverse of average actin filament length | $\approx 5$ $\mu$m$^{-1}$ |
| $q$ | inverse of length over which resistance increases | $\approx 1.7$ $\mu$m$^{-1}$ |
| $\alpha$ | mechanical resistance parameter | $\approx 0.075$ |
| $\kappa_1$ | actin disassembly rate at the front | $\approx 1.25$ min$^{-1}$ |
| $\kappa_2$ | actin disassembly rate at the rear | $\approx 0.80$ min$^{-1}$ |
| $\nu_1$ | Arp2/3-mediated actin assembly rate at the front | $\approx 1.9$ min$^{-1}$ |
| $\nu_2$ | Arp2/3-mediated actin assembly rate at the rear | $\approx 0.90$ min$^{-1}$ |
| $\xi_1$ | detachment rate of Arp2/3 at the front | $\approx 0$ min$^{-1}$ |
| $\xi_2$ | detachment rate of Arp2/3 at the rear | $\approx 1.5$ min$^{-1}$ |
| $\eta_1$ | actin-dependent Arp2/3 attachment rate at the front | $\approx 0$ min$^{-1}$ |
| $\eta_2$ | actin-dependent Arp2/3 attachment rate at the rear | $\approx 0.6$ min$^{-1}$ |

#### PHASE-SPACE PORTRAITS FOR F-ACTIN-ARP2/3-VELOCITY SYSTEM

To extend the phase-space analysis shown in Fig. 1C in the main text, we computed direction fields for some pairwise combinations (other than those shown in Fig. 1C in the main text) of the five variables: edge velocity ($v$), F-actin ($a_1$, $a_2$) and Arp2/3 ($r_1$, $r_2$) in the leading and trailing layers. These fields were computed by binning the observed values of each variable pair, then averaging the observed temporal increments (first differences) within each bin. The resulting direction fields are shown in Fig. S1. Each plot represents a 2D projection of the underlying multidimensional dynamics. The consistent presence of spiraling trajectories suggests that the lamellipodial dynamics is characterized by damped oscillations. Three-dimensional ($r_1 - a_1 - v$) phase field is shown in Fig. S2 revealing the stable steady state at the origin.

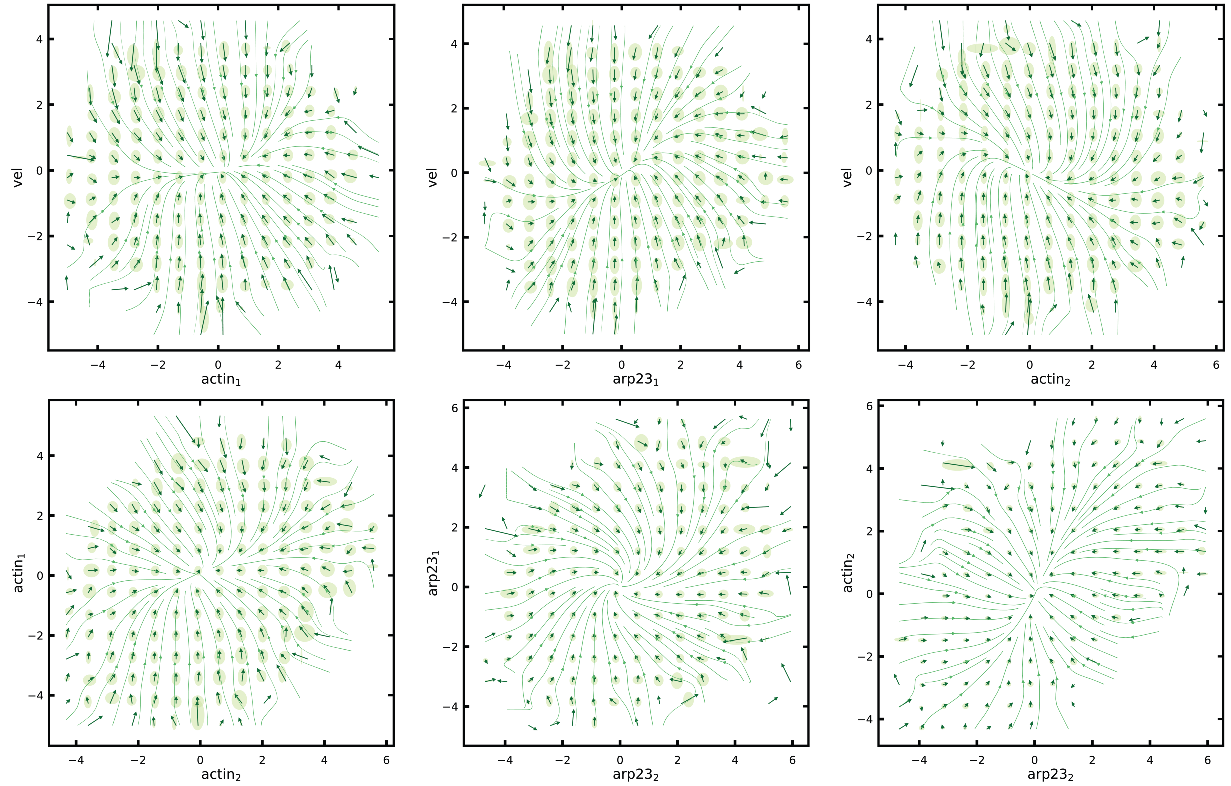


Fig. S1. **Phase-space direction fields for pairs of velocity, actin, and Arp2/3 variables.** Each panel shows a vector field estimated from the binned phase space of two variables, derived from time-differenced experimental data across all cells, spatial compartments and time points. Arrows denote the average direction of variables' change for each bin, and the overlaid green ellipses represent standard deviations of the displacement vectors. The variables include protrusion velocity, actin ($a_1$, $a_2$) and Arp2/3 ($r_1$, $r_2$) intensities in both leading and trailing lamellipodial layers. The data for the averaging is from the total of 972 windows from all 12 cells, with 301-point time series per window (total duration 15 min, with $dt = 3$ sec).

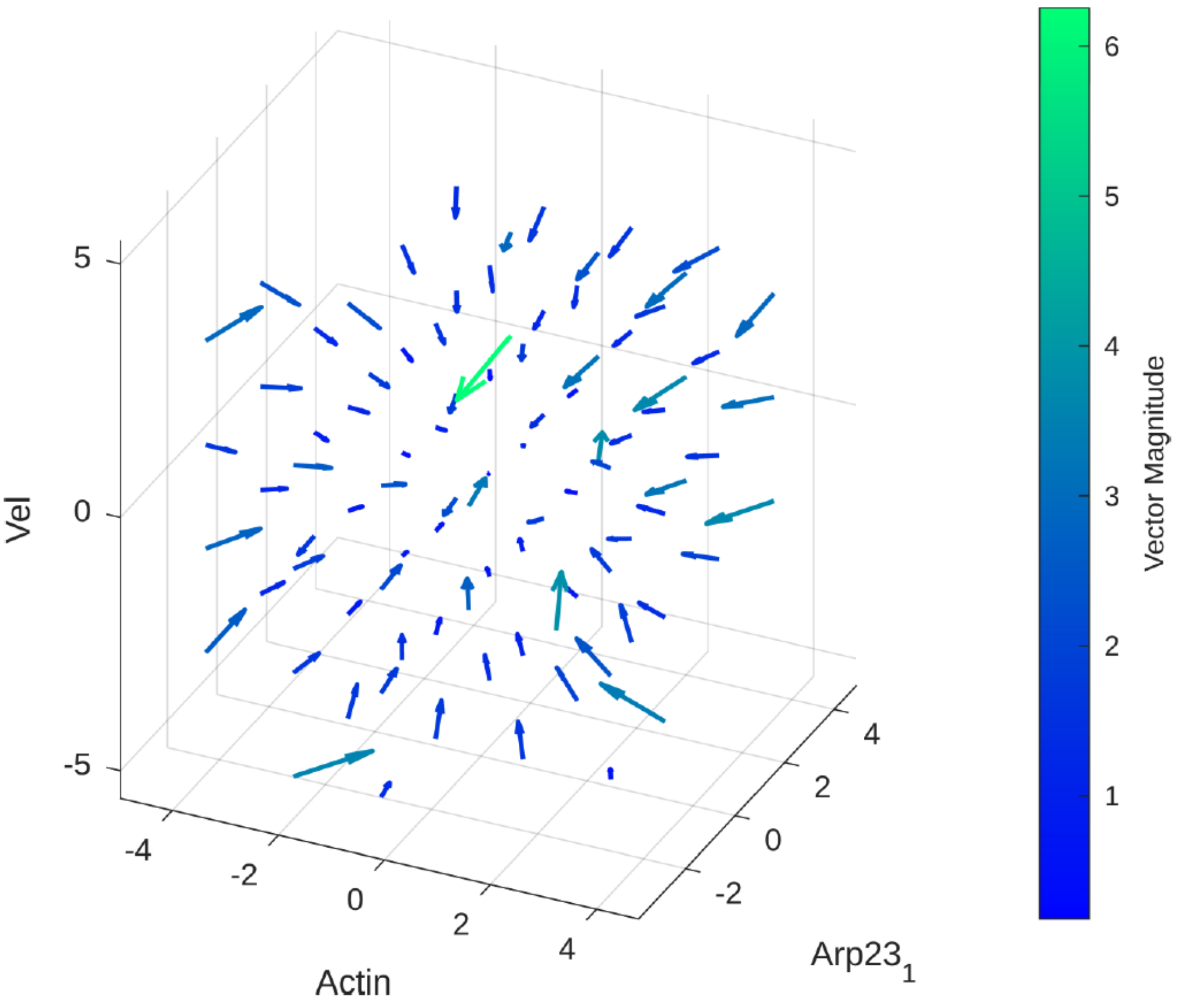


Fig. S2. **Phase-space structure reveals globally coupled oscillations in the actin–Arp2/3–velocity system.** Three-dimensional direction field in the ($Arp2/3_1$, $Actin_1$, Velocity) space. Each arrow denotes the average increment vector (with three vector elements being average changes of ($Arp2/3_1$, $Actin_1$, Velocity) over $dt = 3$ sec), with color and length encoding vector magnitude. The data for the averaging is from the total of 972 windows from all 12 cells, with 301-point time series per window (total duration 15 min, with $dt = 3$ sec).

#### SPARSITY-BASED MODEL SELECTION USING LASSO REGRESSION

To reconstruct an ODE system for F-actin, Arp2/3, and edge velocity, we initially applied ordinary least squares regression without regularization which resulted in the equations shown in Fig. S3A. To enforce sparsity and make sure all terms in this system are meaningful, we employed LASSO (Least Absolute Shrinkage and Selection Operator) regression, which assigns weight to sparsity and penalizes coefficients with low L1 norm. Specifically, let $N$ be the number of examined data points $y$, let $x = (x_1, x_2, \ldots, x_p)$ be the variables vector (i.e., velocity, F-actin, Arp2/3), let $\beta = (\beta_1, \beta_2, \ldots, \beta_p)$ be the coefficients vector, let $X$ be the variables matrix, so that $X_{ij} = (x_i)_j$, and let $\lambda$ be the regularization parameter. LASSO procedure is to find the set of parameters minimizing the following expression:

$$\min_{\beta \in \mathbb{R}^p} \left\{ \frac{1}{N} \|y - X\beta\|_2^2 + \lambda \|\beta\|_1 \right\}.$$

The greater the regularization parameter $\lambda$ is, the smaller is the number of non-zero system parameters.

We gradually increasing parameter $\lambda$ until all coefficients were eliminated. Variables that vanish at low $\lambda$ are less essential, while those that persist at higher $\lambda$ contribute more substantially to the model. Fig. S3B-D shows the regression coefficients as functions of $\lambda$ for each equation. Vertical dashed lines mark the threshold where each term vanishes. LASSO analysis suggests that all terms in each of three equations contribute nontrivially, as all terms vanish only at large values of $\lambda > 1$. Even the relatively small coefficient of the term proportional to the F-actin density in the equation for Arp2/3 vanishes only at $\lambda \approx 4.3$. This suggests that all terms are meaningful.

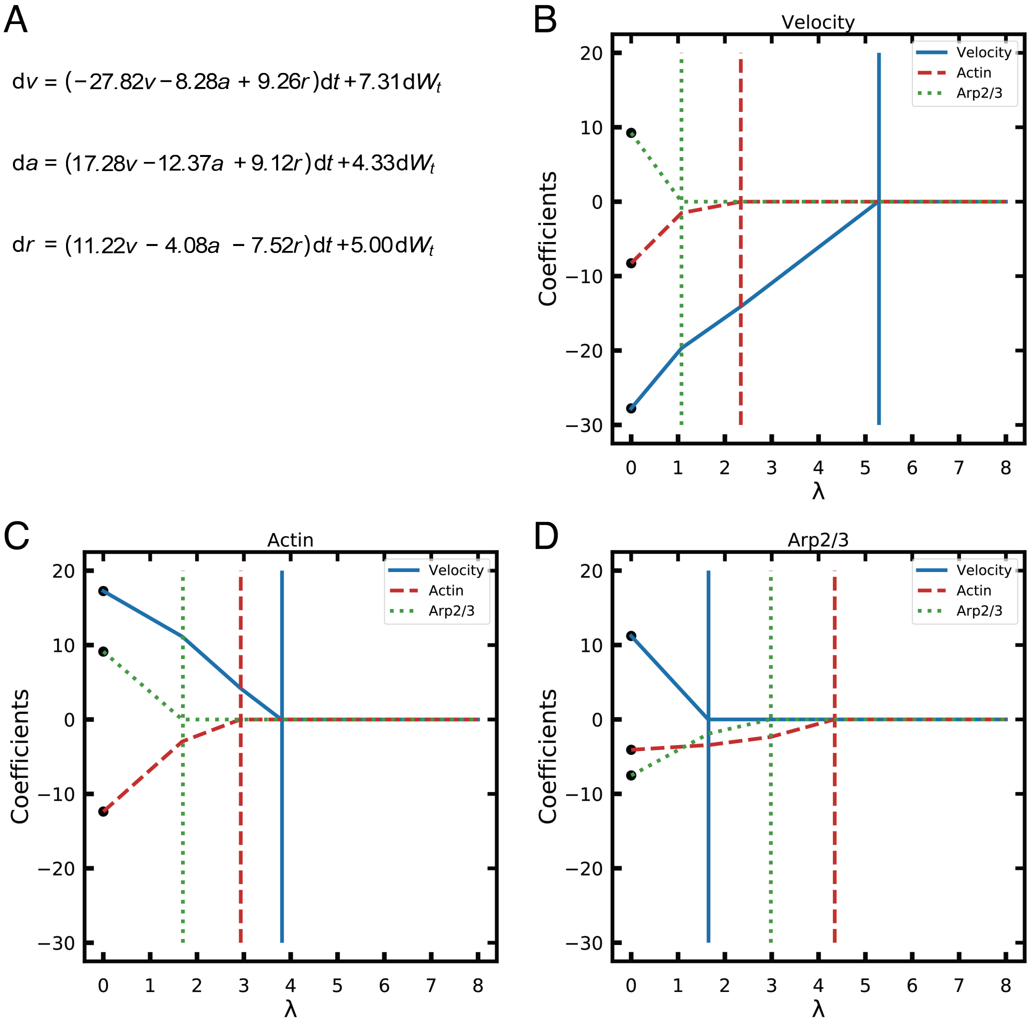


Fig. S3. **LASSO-based analysis of the model.** (A) Ordinary least squares regression result. (B-D) LASSO regression coefficients as functions of the regularization parameter $\lambda$ for each of the three equations. Each curve represents the coefficient associated with a variable, color-coded and line-styled by variable identity (velocity: blue, solid; F − actin: red, dashed; Arp2/3: green, dotted. Vertical lines mark the critical values of $\lambda$ at which the corresponding coefficient vanishes. The regressions are based on the data from the total of 972 windows from all 12 cells, with 301-point time series per window (total duration 15 min, with $dt = 3$ sec).

#### NONLINEAR TERMS CAN BE OMITTED IN THE RECONSTRUCTED ODE MODEL

To examine whether nonlinear terms improve predictive accuracy of the linear regression for the ODE model, we included quadratic terms into the regression corresponding to a second-order perturbation expansion. The resulting fitted system is shown below, with nonlinear terms highlighted in blue:

$$\mathrm{d}v = \left(-23.38v - 2.95a_1 + 18.34r_1 - 10.15a_2 + 5.35\frac{\partial^2 v}{\partial y^2}\right)\mathrm{d}t + 6.75\mathrm{d}W_t^v$$

$$\mathrm{d}a_1 = \left(7.22v - 12.43a_1 + 9.38r_1 - 0.97va_1 + 1.80\frac{\partial^2 a_1}{\partial y^2}\right)\mathrm{d}t + 4.33\mathrm{d}W_t^{a_1}$$

$$\mathrm{d}r_1 = \left(1.30v - 5.80a_1 - 6.10r_1 - 1.08vr_1 + 1.80\frac{\partial^2 r_1}{\partial y^2}\right)\mathrm{d}t + 5.02\mathrm{d}W_t^{r_1}$$

$$\mathrm{d}a_2 = \left(10.56v + 4.08a_1 - 9.37a_2 + 4.30r_2 + 3.67v(a_1 - a_2) + 1.80\frac{\partial^2 a_2}{\partial y^2}\right)\mathrm{d}t + 3.15\mathrm{d}W_t^{a_2}$$

$$\mathrm{d}r_2 = \left(7.72v + 9.42r_1 + 3.11a_2 - 13.87r_2 + 3.43v(r_1 - r_2) + 1.80\frac{\partial^2 r_2}{\partial y^2}\right)\mathrm{d}t + 4.46\mathrm{d}W_t^{r_2}$$

We observe that the nonlinear terms are close to unity (considering that factors $(a_1 - a_2)$ and $(r_1 - r_2)$ are mostly smaller than 1), only comparable to the smallest linear term $1.30v$, and smaller compared to the next smallest linear terms $\sim 3$, not to mention the dominant linear terms $\sim 10$. This suggests that nonlinear terms in the regression may not be meaningful. For the reasons of model sparsity and interpretability, we therefore opted to exclude the nonlinear terms from the reconstructed parsimonious ODE model.

**MEMORY TERMS DO NOT YIELD INTERPRETABLE DYNAMICS**

We also tested a linear model with memory to explore whether including time-delay terms would improve the reconstruction. This is analogous to approaches used in modeling viscoelasticity, where past variables' values influence current dynamics. Rather than incorporating all possible lagged cross-terms, we restricted memory to autoregressive terms, that is, the equation for each variable includes only delay terms proportional to the same variable from the previous four time steps, in addition to its current value. The fitted equations are shown below. Here, for brevity, $(t-i)$ means $(t-i\Delta t)$. Memory terms are highlighted in blue:

$$\mathrm{d}v^t = \left(-25.53v^t + 4.69v^{t-1} + 0.93v^{t-2} - 1.52v^{t-3} - 4.57v^{t-4} - 1.98a_1 + 17.73r_1 - 9.17a_2 + 5.51\frac{\partial^2 v}{\partial y^2}\right)\mathrm{d}t + 6.75W_t^v,$$

$$\mathrm{d}a_1^t = \left(6.72v + 11.68a_1^t - 18.63a_1^{t-1} - 4.14a_1^{t-2} - 0.56a_1^{t-3} - 2.17a_1^{t-4} + 6.71r_1 + 0.49\frac{\partial^2 a_1}{\partial y^2}\right)\mathrm{d}t + 4.33W_t^{a_1},$$

$$\mathrm{d}r_1 = \left(2.98v - 0.93a_1 + 10.17r_1^t - 16.61r_1^{t-1} - 4.12r_1^{t-2} - 0.43r_1^{t-3} - 2.98r_1^{t-4} + 0.49\frac{\partial^2 r_1}{\partial y^2}\right)\mathrm{d}t + 5.02\mathrm{d}W_t^{r_1},$$

$$\mathrm{d}a_2 = \left(5.19v + 3.89a_1 + 13.89a_2^t - 8.99a_2^{t-1} - 10.39a_2^{t-2} - 2.56a_2^{t-3} - 2.88a_2^{t-4} + 3.42r_2 + 0.49\frac{\partial^2 a_2}{\partial y^2}\right)\mathrm{d}t + 3.15\mathrm{d}W_t^{a_2},$$

$$\mathrm{d}r_2 = \left(4.55v + 8.43r_1 + 6.96a_2 + 2.35r_2^t - 12.97r_2^{t-1} - 4.13r_2^{t-2} - 1.64r_2^{t-3} - 3.12r_2^{t-4} + 0.49\frac{\partial^2 r_2}{\partial y^2}\right)\mathrm{d}t + 4.46\mathrm{d}W_t^{r_2}.$$

These results reveal several inconsistencies. First, in the velocity equation, one would expect the coefficients in front of the past velocity values to decay with time, reflecting diminishing influence at longer delays. Instead, the memory coefficients fluctuate in both sign and magnitude, with no interpretable decay pattern. Second, for all molecular species ($a_1$, $r_1$, $a_2$, $r_2$), the inclusion of memory terms reverses the sign of the coefficient (from negative to positive) in front of the variable at the current time, while the lagged coefficients vary irregularly. This not only violates typical expectations of smooth, thin-tailed memory kernels but also obscures physical interpretation. Moreover, adding memory terms affects the magnitude of other variables' coefficients, suggesting that the added complexity dilutes rather than refines the model's explanatory structure. Due to these issues, we conclude that memory terms introduce instability and obscure mechanistic clarity. They neither enhance predictive value nor align with known biochemical feedback pathways. We therefore exclude them from the linear model.

It is worth noting the conceptual distinction from Granger-causal inference methods [1], which include temporal history in a different way. In Granger analysis, the current value of a variable is regressed directly onto its own past and the past of other variables. For example,

$$X^t = c_1X^{t-1} + c_2X^{t-2} + c_3X^{t-3} + c_4X^{t-4} + a_1Y^{t-1} + a_2Y^{t-2}$$

where $X$ and $Y$ are two independent variables, $a$ and $c$ are coefficients. By contrast, our modeling framework is dynamical: we model the time derivative $\frac{dX}{dt}$ as a function of past states $X_{t-j}$. While both approaches incorporate memory, the interpretation and implementation are distinct. Our model aims to reflect continuous-time biophysical dynamics, whereas Granger inference focuses on predictive statistical dependence.

#### SINDY IS NOT SUITABLE FOR OUR SYSTEM DUE TO THE NOISY DATA IN THE VICINITY OF THE SINGLE STABLE STEADY STATE

To evaluate whether existing automated sparse regression methods can accurately recover the linear model equations, we tested the SINDy (Sparse Identification of Nonlinear Dynamical Systems) framework [5]. SINDy uses sparse regression to infer a dynamical system from time series data.

We applied SINDy to mock data generated from our regression-based model. The system used for data generation was an SDE (stochastic differential equation) system with known coefficients and added white noise:

$$\begin{aligned}
\mathrm{d}v &= (-23.39v - 2.94a_1 + 18.33r_1 - 10.15a_2)\,\mathrm{d}t + 6.75\mathrm{d}W_t^{v}\\
\mathrm{d}a_1 &= (7.18v - 12.44a_1 + 9.40r_1)\,\mathrm{d}t + 4.33\mathrm{d}W_t^{a_1}\\
\mathrm{d}r_1 &= (1.30v - 5.79a_1 - 6.13r_1)\,\mathrm{d}t + 5.02\mathrm{d}W_t^{r_1}\\
\mathrm{d}a_2 &= (10.38v + 4.08a_1 - 9.57a_2 + 4.40r_2)\,\mathrm{d}t + 3.15\mathrm{d}W_t^{a_2}\\
\mathrm{d}r_2 &= (7.36v + 9.70r_1 + 2.95a_2 - 13.91r_2)\,\mathrm{d}t + 4.46\mathrm{d}W_t^{r_2}
\end{aligned}$$

SINDy was applied to the resulting mock trajectories using a linear function library. Its output was:

$$\begin{aligned}
\frac{\mathrm{d}v}{\mathrm{d}t} &= -0.20v - 4.17a_1 + 8.52r_1 - 7.75a_2 - 1.65r_2\\
\frac{\mathrm{d}a_1}{\mathrm{d}t} &= 3.31v - 2.88a_1 + 8.73r_1 - 3.06a_2 - 1.11r_2\\
\frac{\mathrm{d}r_1}{\mathrm{d}t} &= -3.41v - 8.30a_1 + 7.64r_1 + 1.31a_2 - 4.01r_2\\
\frac{\mathrm{d}a_2}{\mathrm{d}t} &= 10.15v + 2.31a_1 + 0.63r_1 - 4.55a_2 + 2.15r_2\\
\frac{\mathrm{d}r_2}{\mathrm{d}t} &= 6.36v - 1.10a_1 + 5.46r_1 - 4.88a_2 + 0.25r_2
\end{aligned}$$

Although SINDy correctly predicted the signs of many terms, it failed to recover accurate coefficients. Notably, (i) the self-interaction terms, which dominate the stability and timescales of the system, were severely underestimated. For instance, the velocity coefficient in the velocity equation changed from $-23.39$ (our model) to $-0.20$ (SINDy). (ii) The coefficient in front of $a_1$ in the equation for $a_1$ was reduced from $-12.44$ to $-2.88$, and the coefficient in front of $r_2$ in the equation for $r_2$ – from $-13.91$ to 0.25. (iii) Several signs flipped compared to the ground truth. In particular, the coefficient in front of $r_1$ in the equation for $r_1$ changes from $-6.13$ to 7.64, which reverses the sign of the predicted feedback. Altogether, mean absolute error of coefficients' values was 4.86, and the number of sign mismatches was 4 out of 18.

The key reason for these discrepancies is that SINDy is designed for problems characterized by transient deterministic trajectories with limited noise, while our data consist of small relaxations to the stable steady state after a significant noise 'kicks' the system out of the equilibrium. This dominating noise perturbs the derivative estimates and causes regularization-based methods to misattribute variability to incorrect features or to shrink dominant coefficients.

#### CELL-TO-CELL HETEROGENEITY DOES NOT SIGNIFICANTLY AFFECT INFERRED DYNAMICS

Biological systems are inherently heterogeneous, and single-cell variability can potentially influence the outcome of any regression-based inference. In our study, the main text reports results from the pooled dataset comprising fluorescence time series of velocity, actin, and Arp2/3 from 12 individual cells. A natural concern is whether differences between individual cells might lead to significantly different inferred dynamics or even qualitatively distinct mechanistic models.

To evaluate this possibility, we applied our regression pipeline separately to each of the 12 cells. The resulting coefficients for each dynamical equation are shown in Fig. S4. In each subplot: (i) Black symbols represent the coefficients estimated from individual cells; (ii) The green solid line denotes the coefficient obtained from the full dataset pooled across all cells; (iii) The blue dashed line is the mean of the 12 single-cell regressions; (iv) The blue shaded region indicates $\pm$ one standard deviation across 12 cells.

As shown, the majority of single-cell coefficients cluster closely around the pooled estimate and lie well within one standard deviation of the mean. In particular, all larger terms remain consistent across cells. This suggests that although cells exhibit intrinsic variability, the underlying dynamical architecture governing F-actin and Arp2/3 regulation and feedbacks between velocity and molecular densities is robust and reproducible. Therefore, we conclude that inter-cell heterogeneity does not significantly compromise the conclusions drawn from the pooled model. The dominant features of the inferred dynamics are conserved across cells, justifying the use of the population-averaged regression.

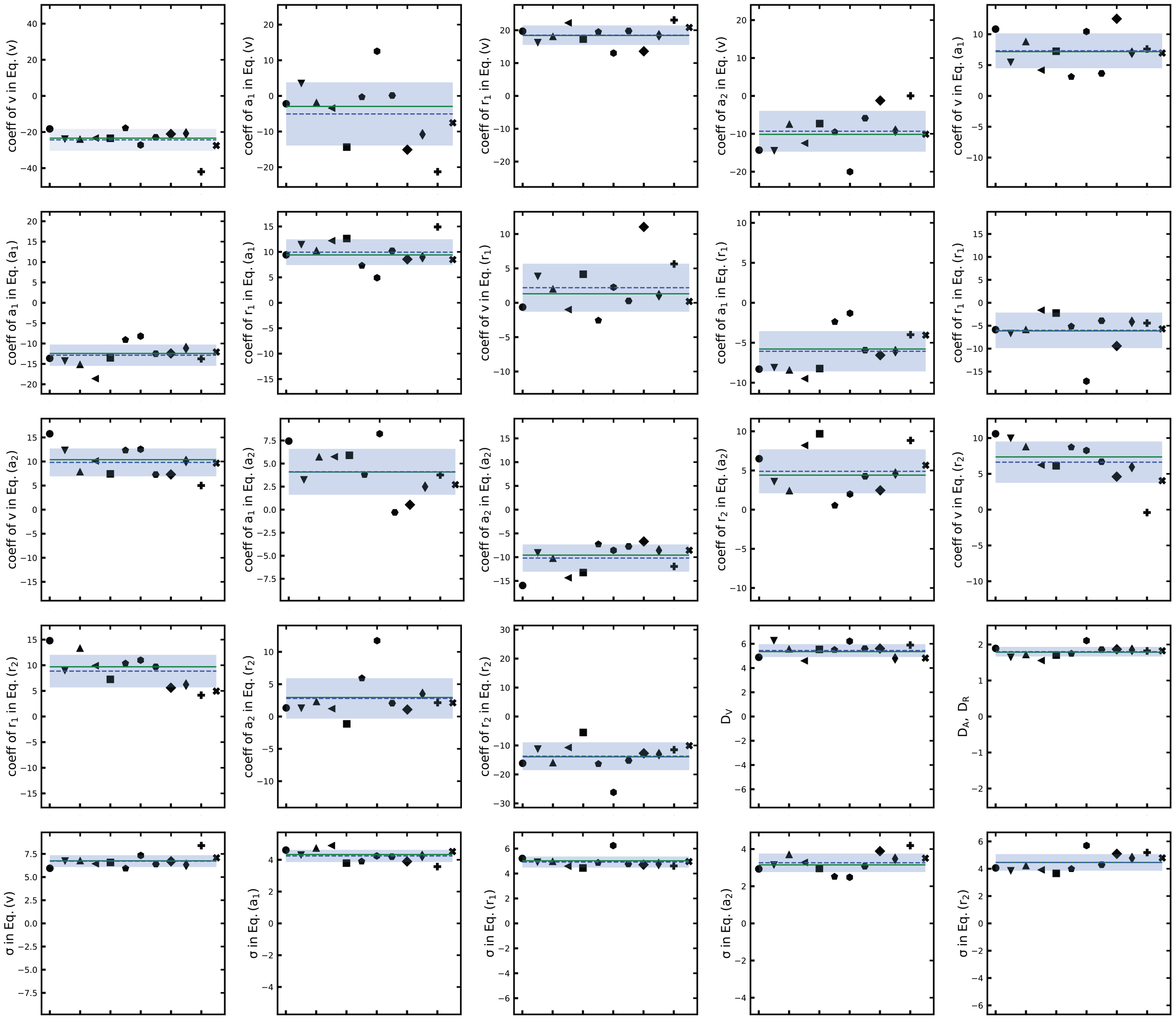


Fig. S4. **Single-cell regressions show consistent dynamics across heterogeneous cells.** Each subplot shows the value of one regression coefficient across the 12 cells (black symbols). The green solid line indicates the value estimated from the pooled dataset, as used in the main text. The blue dashed line shows the mean of the single-cell estimates, and the blue shaded region spans ± one standard deviation.Results from the pooled data are from the total of 972 windows from all 12 cells, Results from the individual cells are from 81 windows each. Each window provides 301-point time series (total duration 15 min, with $dt = 3$ sec).

#### 2D PROBABILITY DISTRIBUTIONS

To complement the subset of joint probability distributions shown in the main text, we computed all pairwise 2D distributions among the five key variables: edge velocity ($v$), actin ($a_1$, $a_2$) and Arp2/3 ($r_1$, $r_2$) in the leading and trailing compartments. The results that are not shown in the main text are displayed in Fig. S5, where blue denotes experimental data and orange indicates model simulations.

Each subplot shows a 2D scatter plot with marginal 1D distributions along the axes. The elliptical structures of the scatter plots indicate that all pairwise combinations of variables are approximately jointly Gaussian. The orientations and elongations of the ellipses reflect the correlation strength and direction between the respective variables. Marginal 1D distributions also consistently reveal unimodal, symmetric distributions, further supporting the assumption of Gaussian white noise underlying the stochastic model.

Importantly, the simulated data (orange) accurately accounts for both the shape and spread of the experimental distributions (blue), indicating that the inferred model captures not only marginal variability but also pairwise statistical dependencies among variables across the spatial layers.

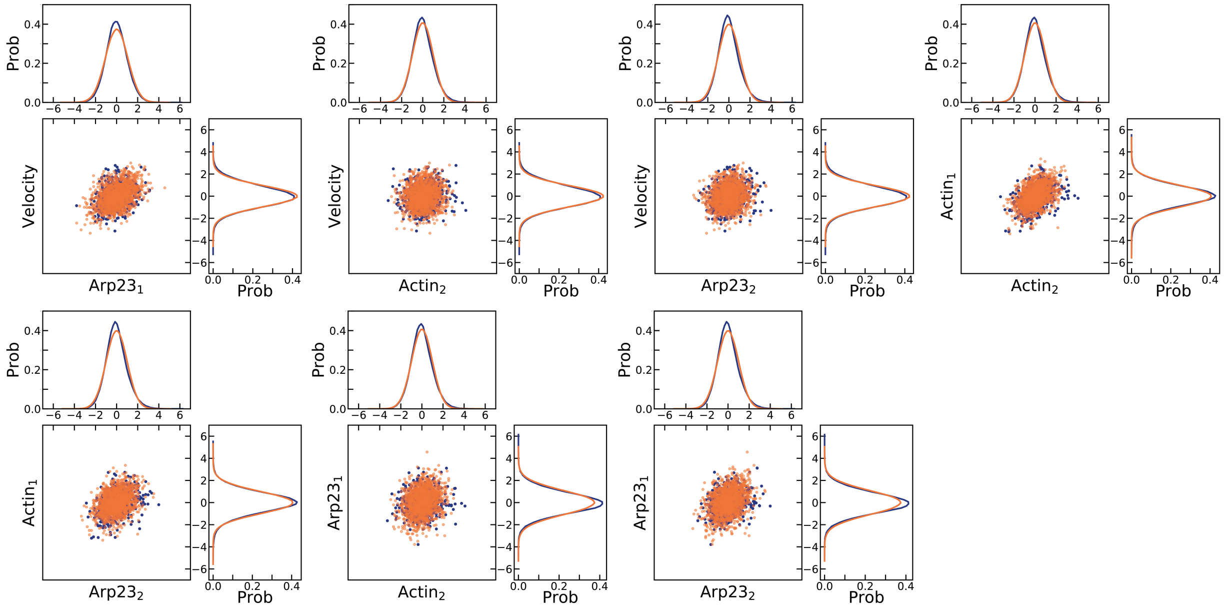


Fig. S5. **Joint 2D distributions between all pairs of variables.** Scatter plots and marginal 1D distributions comparing experimental data (blue) and model simulations (orange) for all pairs of variables (velocity ($v$), actin ($a_1$, $a_2$), and Arp2/3 ($r_1$, $r_2$)). Each central panel shows the 2D distribution of paired variables, while the top and side panels display their respective 1D marginals. The data for the averaging is from the total of 972 windows from all 12 cells, with 301-point time series per window (total duration 15 min, with $dt = 3$ sec).

#### CROSS-CORRELATION ANALYSIS

To test the model, we computed cross-correlations between all pairs of variables. The complete set of lagged correlation curves is presented in Fig. S6, where blue denotes cross-correlations obtained from the experimental data, and orange denotes cross-correlations from simulations of the inferred stochastic model. The quantitative agreement between experiment and simulation across these metrics validates the model. (The only small disagreement is that the data shows almost instant correlation between F-actin and Arp2/3 in the leading compartment, while the model predicts that F-actin increase follows Arp2/3 increase at a slightly earlier time.)

The analysis reveals that, first, the relationship between edge velocity and Arp2/3 density at the edge is characterized by rapid synchronization. The cross-correlation between $r_1$ and velocity exhibits a sharp peak at zero lag ($\tau \approx 0$). This indicates that fluctuations in protrusion velocity are immediately coupled to Arp2/3 density at the leading edge. This observation supports the mechanistic assumption that Arp2/3 incorporation is directly driven by instantaneous velocity rather than accumulating slowly and that uncapped Arp2/3-anchored filaments drive the protrusion.

Second, the interaction between F-actin and velocity reveals a distinct feedback loop. As seen in the velocity-actin couplings (Fig. 2B in the main text, left and Fig. S6), a negative correlation peak occurs at negative lag ($\tau < 0$), confirming that a higher F-actin density decelerates edge velocity (according to our hypothesis, due to mechanical resistance). Conversely, a positive correlation emerges at positive lag ($\tau > 0$), reflecting that higher protrusion velocity leads to F-actin accumulation at the edge due to polymerization, and at the rear due to advection. Third, Arp2/3 and F-actin, as well as F-actin globally, display instant correlations.

Collectively, these data is consistent with a mechanical cycle in which velocity and Arp2/3 at the edge are tightly synchronized, while actin accumulates with a delay to provide resistance, all the while being continuously transported away from the leading edge.

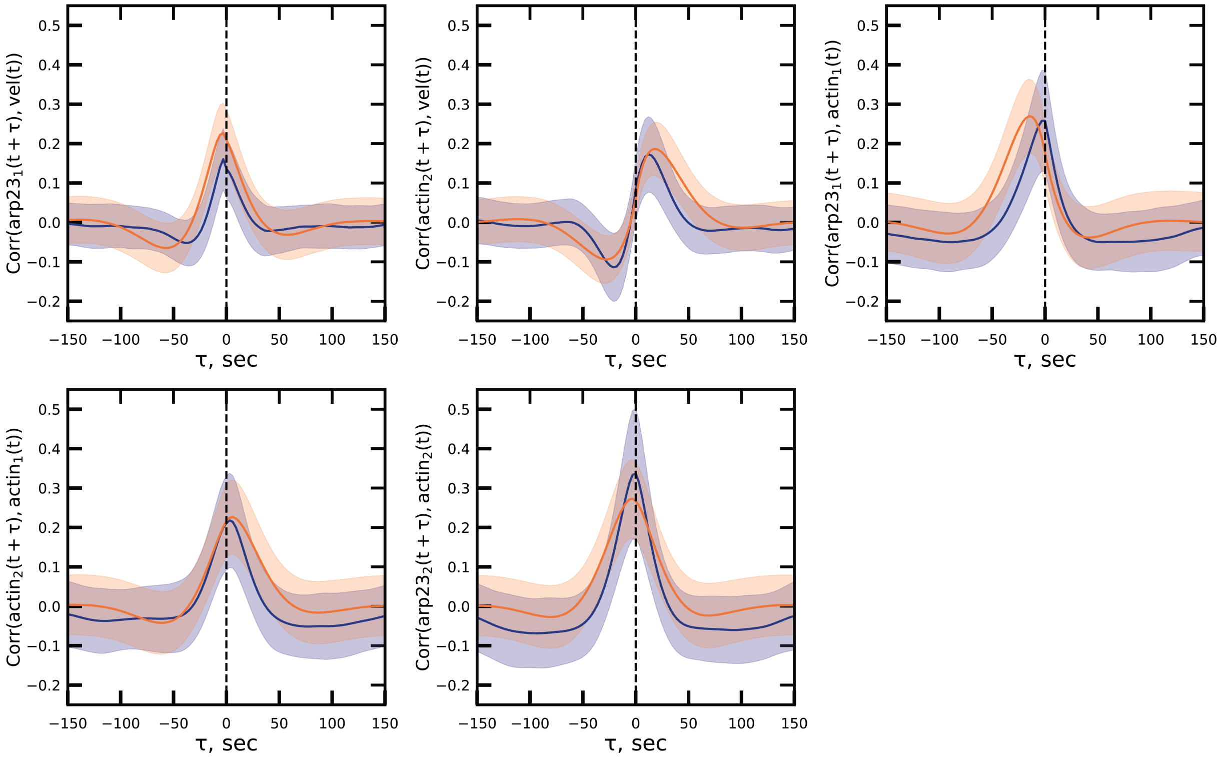


Fig. S6. **Cross-correlation functions between all variable pairs.** Lagged cross-correlation functions between all pairs of actin, Arp2/3, and velocity variables. Blue curves represent experimental measurements; orange curves represent model simulations. Shaded regions indicate $\pm$ standard deviation. The data for the averaging is from the total of 972 windows from all 12 cells, with 301-point time series per window (total duration 15 min, with $dt = 3$ sec).

#### UPSTREAM WAVE DYNAMICS COULD AMPLIFY ARP2/3 NOISE AND DRIVE STOCHASTIC EDGE BEHAVIOR

We hypothesized that the discrepancy between the simple Poisson process of the Arp2/3 and F-actin molecular noise and the observed higher level of respective noise (Fig. S7B) can be explained by stochastic fluctuations of a relatively small number of WAVE molecules activating Arp2/3 at the edge. To test this hypothesis, we first simulated a simplified stochastic discrete model in which upstream fluctuations of WAVE numbers propagate downstream. In this model (Fig. S7A), WAVE molecules attach and detach to the edge with rates $B_{\mathrm{WAVE}}$ and $\xi_{\mathrm{WAVE}}$ respectively, while Arp2/3 molecules are inserted into the actin network by WAVE molecules with rate $B_R$. After insertion, Arp2/3 molecules drift rearward and dissociate from the network with rate $\xi$.

We simulated this model and varied rates $B_{\mathrm{WAVE}}$ and $\xi_{\mathrm{WAVE}}$ while holding their ratio constant, so that the mean number of WAVE molecules stayed fixed at 10, corresponding to the noise level for WAVE of ∼30%. As shown in Fig. S7D, the resulting noise of the Arp2/3 numbers depends on the WAVE turnover ratio $\xi_{\mathrm{WAVE}}/\xi$. When WAVE turns over slowly, its fluctuations are fully inherited by Arp2/3, which also exhibits ∼30% noise. However, when WAVE turns over rapidly, its fluctuations are averaged out in time before Arp2/3 turns over and Arp2/3 noise approaches the Poisson limit of ∼5%. This result demonstrates that a small number of upstream regulators with slow dynamics can be the source of the observed significant lamellipodial noise.

Next, we implemented a stochastic version of the full mechanistic PDE model. In this model (Fig. S7A), all reaction terms, such as actin polymerization, Arp2/3 disassembly, and velocity changes, are treated as probabilistic events. During a small time step $\mathrm{d}t$, an event with rate $k$ occurs with probability $k\mathrm{d}t$. Respective 'birth and death' events for WAVE, Arp2/3 and actin filaments were implemented. The actin dynamics was simplified, so that the explicit polymerization of filaments was neglected; rather, a filament of an average length was branched with an effective rate given by the PDE model parameters, and then deleted at once with another rate due to disassembly. Spatial advection was modeled by deterministic retrograde shifts of Arp2/3 molecules and actin filaments, while diffusion processes were modeled by random lateral shifts.

A snapshot of this stochastic simulation is shown in Fig. S7C. The middle panel shows the instant spatial distribution of Arp2/3 (orange dots) and actin filaments (segments), while the top and side panels show the edge velocity profile and averaged (along the direction parallel to the edge) Arp2/3 concentration, respectively. Despite constant mean parameters, molecular distributions and velocity profiles exhibit significant spatial heterogeneity, driven by the noise in WAVE turnover, stochastic events of Arp2/3 recruitment and birth/death of actin filaments. This simulation captured the experimentally observed variability in both molecular concentrations and edge velocity, providing slow WAVE turnover, so that the fluctuations of WAVE numbers are not averaged out during the characteristic time of the Arp2/3-actin treadmill. Though we cannot exclude the possibility of significant experimental measurement noise, which could account for a large part of the observed fluctuations, these results demonstrate that large fluctuations of Arp2/3, actin and velocity could arise from slow turnover of small number of WAVE activators of Arp2/3.

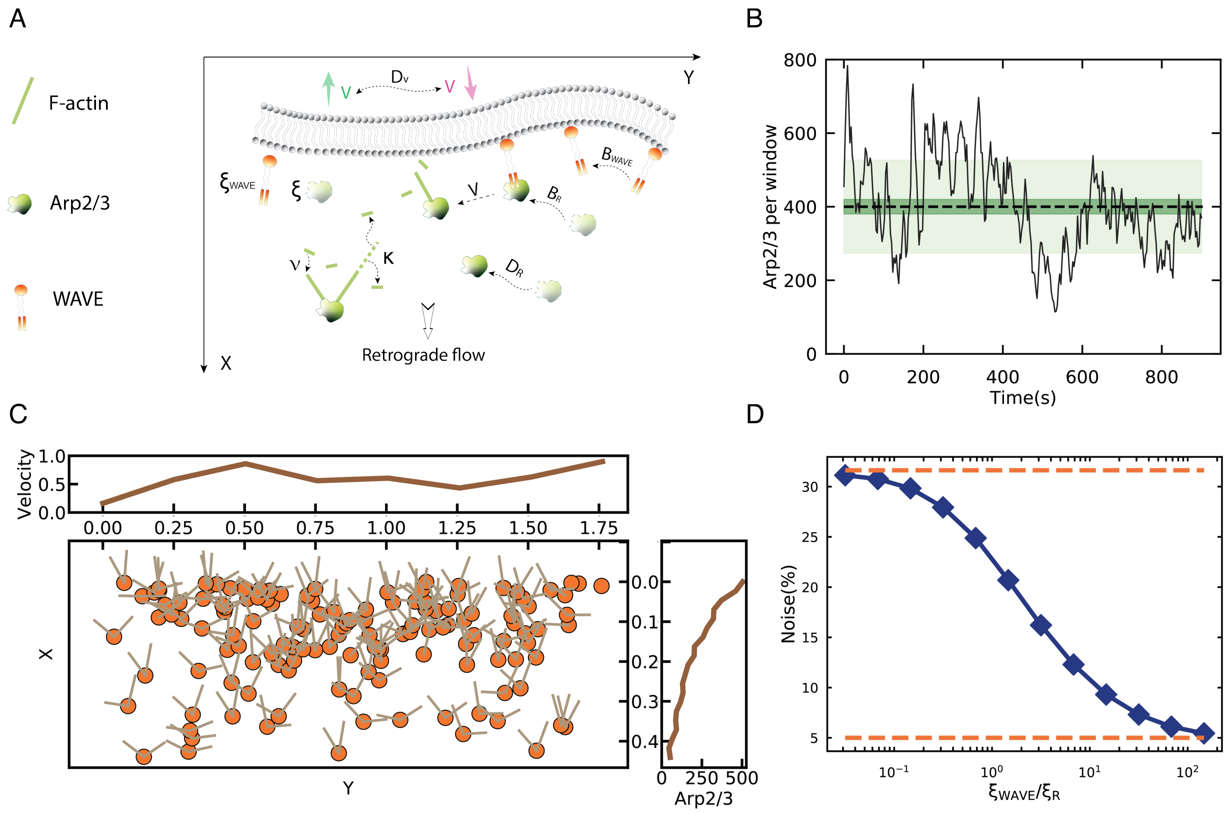


Fig. S7. **Stochastic modeling reveals noise amplification in Arp2/3 dynamics driven by upstream WAVE fluctuations.** (A) Schematic of the model. Arp2/3 attachment to the F-actin network is controlled by WAVE activation. WAVE molecules appear at the edge at rate $B_{\mathrm{WAVE}}$ and degrade at rate $\xi_{\mathrm{WAVE}}$. In turn, Arp2/3 is generated proportionally to the local WAVE number, with birth rate $B_R$. After appearance, Arp2/3 is activated at the membrane and released into the lamellipodium with velocity $V$. Actin filaments are generated at a rate $\nu$ by Arp2/3 molecules, and disassemble at a rate $\kappa$. Arp2/3 molecules detach at a rate $\xi$. The edge velocity $V$ evolves from the balance between local resistance and polymerization forces and diffuses laterally at a rate $D_v$. (B) Experimental time series of Arp2/3 concentration near the leading edge. The black line demonstrates one example of a time series of Arp2/3. The black dashed line represents the average. The light green band denotes respective standard deviation. The dark green envelope corresponds to the expected standard deviation under a Poisson birth–death model. (C) Snapshot from the stochastic simulation of the mechanistic model (see Fig. 3A in the main text), where birth and death events are implemented using a probabilistic method within small time intervals. The central panel shows spatial distributions of Arp2/3 (orange circles) and F-actin (filaments). The top panel displays the corresponding edge velocity profile along the edge, and the right panel shows the Arp2/3 concentration averaged along the lamellipodial width, from the edge to the rear. (D) Computed ratio of the standard deviation to the mean number of Arp2/3 molecules per window as a function of the ratio between WAVE disassembly rate ($\xi_{\mathrm{WAVE}}$) and Arp2/3 disassembly rate ($\xi$).

#### REGRESSION-INFERRED DIFFUSION COEFFICIENTS VARY WITH COARSER SCALING

In our regression model, we included lateral diffusion terms that capture significant spatial coupling along the cell edge. These terms were discretized using finite-difference approximations across neighboring compartments. We investigates how the inferred effective diffusion coefficients behave under the following scaling procedure: let us lump together pairs of laterally neighboring compartments, consider the net F-actin and Arp2/3 signals from these pairs, repeat the regression analysis based on this scaled data and use the finite difference formula for the diffusion terms with the lumped densities and increased spatial step. Then, lump 4 laterally neighboring compartments, etc.

We found the following resulting equation:

$$\frac{x^{t+1} - x^t}{\Delta t} = c_1 v^t + c_2 a_1^t + c_3 r_1^t + c_4 a_2^t + c_5 r_2^t + \Delta(x_L^t - 2x^t + x_R^t), \tag{S14}$$

where $x$ is a variable that can be $a, r$ or $v$, $x_L^t$ and $x_R^t$ denote this variable in neighboring windows in the lateral directions, and $\Delta$ is the transport coefficient. Note that in the data, $x$ is the value of the variable *averaged at each time step* across $N$ windows that are lumped together (or, for $N = 1$, is just the variable in one window at that time step).

We found that the transport coefficients scale approximately as $\Delta \sim 1/\sqrt{N}$ with the number of the lumped windows (Fig. S8). (The reaction terms remained approximately constant with the coarse-graining.) The plausible explanation for this result is as follows.

Let us note that one possible mechanism of the lateral transport of F-actin is the lateral component of the uncapped actin filament growth with a rate on the order of 1 $\mu$m/min (see the main text). Over the time interval $\Delta t = 3$ sec, this corresponds to the spatial shift on the order of 50 nm. Similarly, the lateral component of the retrograde flow, on the order of a fraction of 1 $\mu$m/min, would cause the lateral shift of tens of nm in the same time. These lateral shifts are much smaller than the window size of $\approx 700$ nm.

We introduce the characteristic length $\delta$ along the leading edge, from which the random lateral shifts of the actin network occur over the time interval $\Delta t = 3$ sec. Then, the net 'amount' of the actin network that shifts into the neighboring window over this time will be $\delta x_\delta^t$, where $x_\delta^t$ is the average density of variable $x$ characterizing the network at time $t$ within the interval of size $\delta$ at the edge of the window *from which the shift occurs.* This influx from the neighboring window will increase the *average density in the whole recipient window* by $\delta x_\delta^t / NL$, where $NL$ is the size of $N$ lumped together $L$-compartments. Finally, note that under a crude assumption that random variable $x$ is uncorrelated in space, the standard deviation of $x_\delta^t$ is $\sqrt{NL/\delta}$-fold greater than the standard deviation of $x^t$. Then, after averaging of the data over many time points, the transport term in Eq. (S14) will look like $\sim (\sqrt{\delta/NL})(x_L^t - 2x^t + x_R^t)$.

Such scaling is indeed obtained from the data (Fig. S8). However, this means that the effective diffusion coefficient obtained from the data is not an accurate estimate of the true diffusion coefficient simply because the spatial resolution of the data is not fine enough.

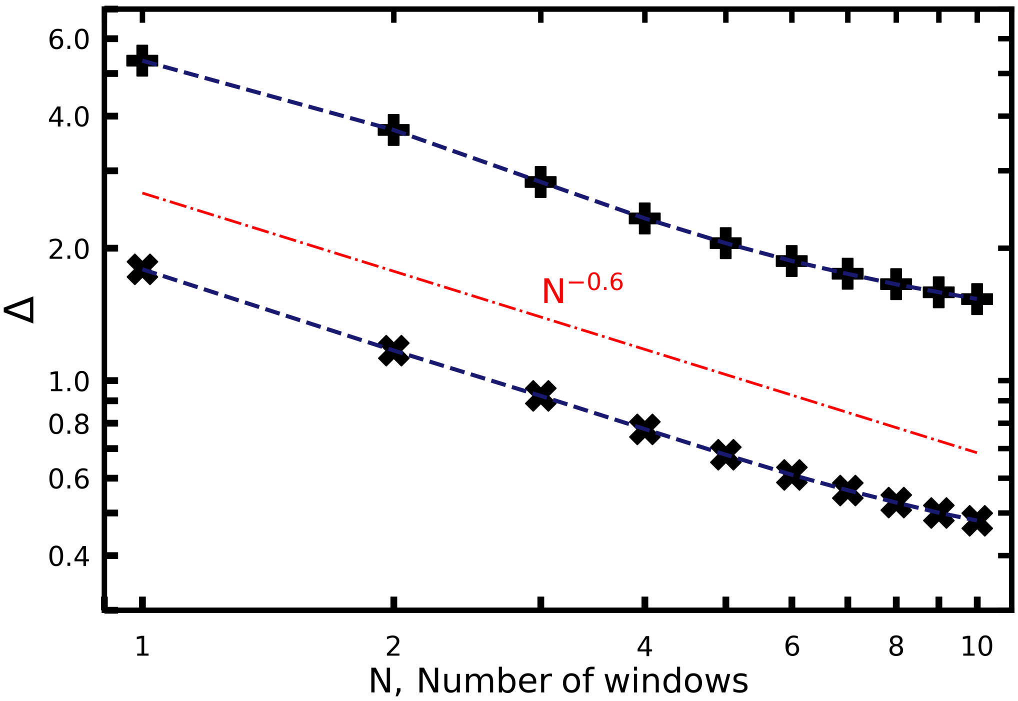


Fig. S8. **Diffusion coefficients scale with the lumped number of neighboring windows.** Estimated transport coefficients (in nondimensional units of $L^2/T$) for edge velocity (marker: plus) and molecular components (e.g., $a_1$, $r_1$, $a_2$, $r_2$; marker: cross) are plotted against the lumped number of neighboring windows used in the regression. Both sets of inferred transport coefficients $\Delta$ exhibit an approximately inverse square root scaling with the increase of the lumped window count. The data for the regression is from the total of 972 windows from all 12 cells, with 301-point time series per window (total duration 15 min, with $dt = 3$ sec).
